# Dynamic, single-cell monitoring of CAR T cell identity and activation with Raman spectroscopy

**DOI:** 10.64898/2026.02.22.707331

**Authors:** Ariel Stiber, Boi Quach, Babatunde Ogunlade, Antony Georgiadis, Kai Chang, Yuanwei Li, Patrick Quinn, Haoqing Wang, Kristin C. Y. Tsui, Charm Ang, Elena Sotillo, David B. Miklos, Crystal Mackall, Zinaida Good, Jennifer A. Dionne

**Affiliations:** Department of Materials Science and Engineering, Stanford University, Stanford, California, USA; Division of Immunology and Rheumatology, Department of Medicine, Stanford University School of Medicine, Stanford, California, USA; Division of Computational Medicine, Department of Medicine, Stanford University School of Medicine, Stanford, California, USA; Center for Cancer Cell Therapy, Stanford Cancer Institute, Stanford University, Stanford, California, USA; Department of Electrical Engineering, Stanford University, Stanford, California, USA; Stanford Cryo-EM Microscopy Center, Stanford University, Stanford, California, USA; Cryoelectron Microscopy, Nucleus at Sarafan ChEM-H, Stanford University, Stanford, California, USA; Division of Hematology and Oncology, Department of Pediatrics, Stanford University, Stanford, CA, USA; Division of Blood and Marrow Transplantation and Cellular Therapy, Department of Medicine, Stanford University, Stanford, California, USA; Parker Institute for Cancer Immunotherapy, Stanford University, Stanford, California, US; Weill Cancer Hub West, Stanford University, Stanford, California, USA; Department of Radiology, Stanford University School of Medicine, Stanford, California, USA

## Abstract

Chimeric antigen receptor (CAR) T cell therapies have reshaped treatment for cancers and immune-mediated diseases, yet their safety and efficacy depend on both the proliferation of engineered cells and their dynamic functional state — features that remain challenging to monitor in real-time clinical settings. Current methods require targeted labels, extensive processing, and provide only static snapshots of cell identity and activation. Here, we introduce a surface-enhanced Raman spectroscopy (SERS) and machine learning (ML) approach that enables single-cell identification of engineered CAR T cells without molecularly-targeted labels and time-resolved, semi-continuous monitoring of their functional activation state through a single physical readout spanning donor-derived cells and patient blood. From intrinsic vibrational signatures of live cells, we detect spectral differences resulting from engineered receptor expression in donor-derived CD19- and GD2-targeted CAR T cells (nine and five donors, respectively) with 81-85% donor-level accuracy, and resolve dynamic antigen-specific activation trajectories with temporal precision. Applying this same SERS-ML method to longitudinal samples from a four-patient CD19-CAR T therapy cohort, we classify patient peripheral blood mononuclear cells across pre-and post-infusion timepoints with a mean patient-level accuracy of 84%, and show that isolated CAR-positive T cells are distinguishable from both CAR-negative T cells and background populations with average 86-88% accuracies. These capabilities stem from biochemical signatures consistent with processes such as receptor expression, tonic signalling, and immune synapse formation, demonstrating a single method that reports both cellular identity and activation state with biochemical specificity across cell engineering and clinical monitoring contexts. Our results extend CAR T cell monitoring beyond static phenotyping and support the potential of SERS-ML analysis for rapid, point-of-care assessment of engineered immune cells across the therapeutic lifecycle.

## Introduction

Chimeric antigen receptor (CAR) T cell therapy has emerged as a transformative immunotherapy with the potential to improve the treatment of cancer, autoimmune, and inflammatory diseases. In this approach, autologous T lymphocytes are isolated and genetically engineered to express CARs, synthetic receptors that direct cytotoxicity towards target cells expressing a chosen antigen. CAR T cell therapies minimize the collateral damage observed with conventional cancer therapies, such as chemotherapy, and elicit more robust immune responses than those typically achieved by antibody-based therapies. CD19-targeted therapies (CD19-CAR) have already changed the standard of care for refractory B cell and plasma cell malignancies^1^ and are expanding into earlier lines of treatment.^2^ Beyond hematologic malignancies, several CAR constructs have shown encouraging clinical activity in solid tumors, including GD2-CAR T cells for diffuse midline gliomas.^3^ Given their growing use and versatility, CAR T cell therapies have become a pillar of modern medicine. Further breakthroughs in this field will require methods that can dynamically monitor engineered cells across the therapeutic lifecycle, from pre-infusion characterization of engineered cell identity and activity to post-infusion tracking of persistence and functional state.

Several obstacles still limit effective CAR T development, particularly the risk of adverse events and variable efficacy. Post-infusion CAR T cell expansion and persistence are important correlates of efficacy and toxicity.^4,5^ Excessive expansion can trigger cytokine release syndrome (CRS) and immune effector cell-associated neurotoxicity syndrome (ICANS), which develop rapidly but can be mitigated if identified early. Conversely, insufficient CAR T cell expansion within the first two weeks post-infusion may indicate treatment failure, necessitating early consideration of alternatives. Further, a co-incidence of fever with a lack of CAR T cell expansion may instead indicate infection, requiring treatment distinct from CRS and ICANS mitigation. Tools to quantify and monitor CAR T cells throughout treatment are therefore vital for both research and clinical care.

Current CAR T cell monitoring methods generally trade off depth of phenotypic cellular information and clinical practicality. Transcriptomic profiling provides detailed molecular information on identity and activation state, but is expensive, slow, labor-intensive, and not routinely performed in clinical workflows.^6^ PCR and flow cytometry are used more commonly; however, PCR does not yield phenotypic information and is not a perfect correlate of circulating cell numbers due to the potential for multiple transgene integration events.^7^ Flow cytometry, on the other hand, is susceptible to batch variability and operator bias, requires pre-planned fluorescent panels and labeling, and can be perturbative to cell function.^8,9^ Additionally, it cannot continuously monitor the phenotypic changes associated with dynamic T cell activation. As a result, CAR T cell monitoring is often time-consuming and expensive, precluding prompt point-of-care support and relegating vital data to retrospective analyses.^10^ Newer label-free techniques, such as ghost cytometry and morphology-based sorting, enable high-throughput cellular phenotyping based on optical properties including size, morphology, and refractive index.^9,11^ However, these methods have not yet been shown to differentiate subtle changes resulting from genetic engineering and, because they primarily capture morphology and optical properties rather than molecular composition, offer limited biochemical specificity. Collectively, these monitoring methods generally offer only static or periodic snapshots of cell states and lack the high temporal resolution and dynamic capabilities needed for real-time analytics.

Here, we present a method for live, single-cell, dynamic CAR T cell quantification and state monitoring without molecularly targeted labels using surface-enhanced Raman spectroscopy (SERS) and machine learning (ML). Raman spectroscopy measures inelastic light scattering from molecular vibrations; the vibrational energies of biomolecules serve as intrinsic labels, avoiding perturbative sample preparation as well as molecular reporters, fluorescent antibodies, or other targeted affinity labels.^12^ In SERS, optically resonant nanostructures amplify local electromagnetic fields under laser illumination, enhancing Raman signals by at least 4 orders of magnitude (and often higher).^13–16^ This increases spatial sensitivity and reduces acquisition times to seconds or sub-seconds. The resulting optical “fingerprints,” when analyzed by an integrated ML algorithm with the ability to recognize patterns in complex datasets, inform identity and real-time responses to various environmental conditions at single-cell resolution. Raman spectroscopy, often combined with ML, has previously been used to classify immune cells^13,17^ and assess activation^18–22^ or drug response,^23^ but has largely focused on chemically stimulated or systemically activated T cells measured at discrete timepoints and has typically compared only static resting versus activated states.

We generated donor-derived Mock (untransduced) and CAR T cells (Fig. 1ai-ii) and analyzed them individually and in co-culture with target cells to study how CAR expression and antigen-specific activation shape Raman spectra (Fig. 1aiii). Using SERS (Fig. 1bi), we obtained spectra with key biomolecular features (Fig. 1bii)^24^ and, through ML analysis of these multivariate signatures (Fig. 1biii), demonstrated both cellular identification and functional monitoring. In this more challenging setting, as compared with distinguishing T cells from unrelated cell types, our technique classified CD19-CAR T cells from Mock with ∼81% average donor-level accuracy across nine donors, and GD2-CAR T cells from Mock with ∼85% accuracy across five donors. Antigen-specific activation in co-culture was distinguished with ∼95% accuracy. Semi-continuous tracking of activation dynamics produced time-resolved spectra that aligned with known activation biomarker progression and flow cytometry trends, demonstrating that Raman spectra can capture dynamic functional states. These results show that both CAR expression and activation state produce detectable spectral shifts, notably in protein-, aromatic-, nucleic acid-, and mitochondrial-associated bands. Finally, we extended our workflow to longitudinal peripheral blood mononuclear cell (PBMC) samples from CD19-CAR T-treated patients, collected before and after infusion. We found that PBMC spectra classified by timepoint with ∼83% average patient-specific accuracy across four patients. Additionally, flow-isolated CAR-positive T cells were distinguishable from CAR-negative T cells and non-CAR background populations with an average ∼84% classification accuracy. Collectively, our method enables dynamic phenotyping of live, donor-derived, and patient-derived CAR T cells without molecularly targeted labels, an important step towards rapid, low-cost, point-of-care monitoring of engineered cell therapies.

**Fig. 1:**
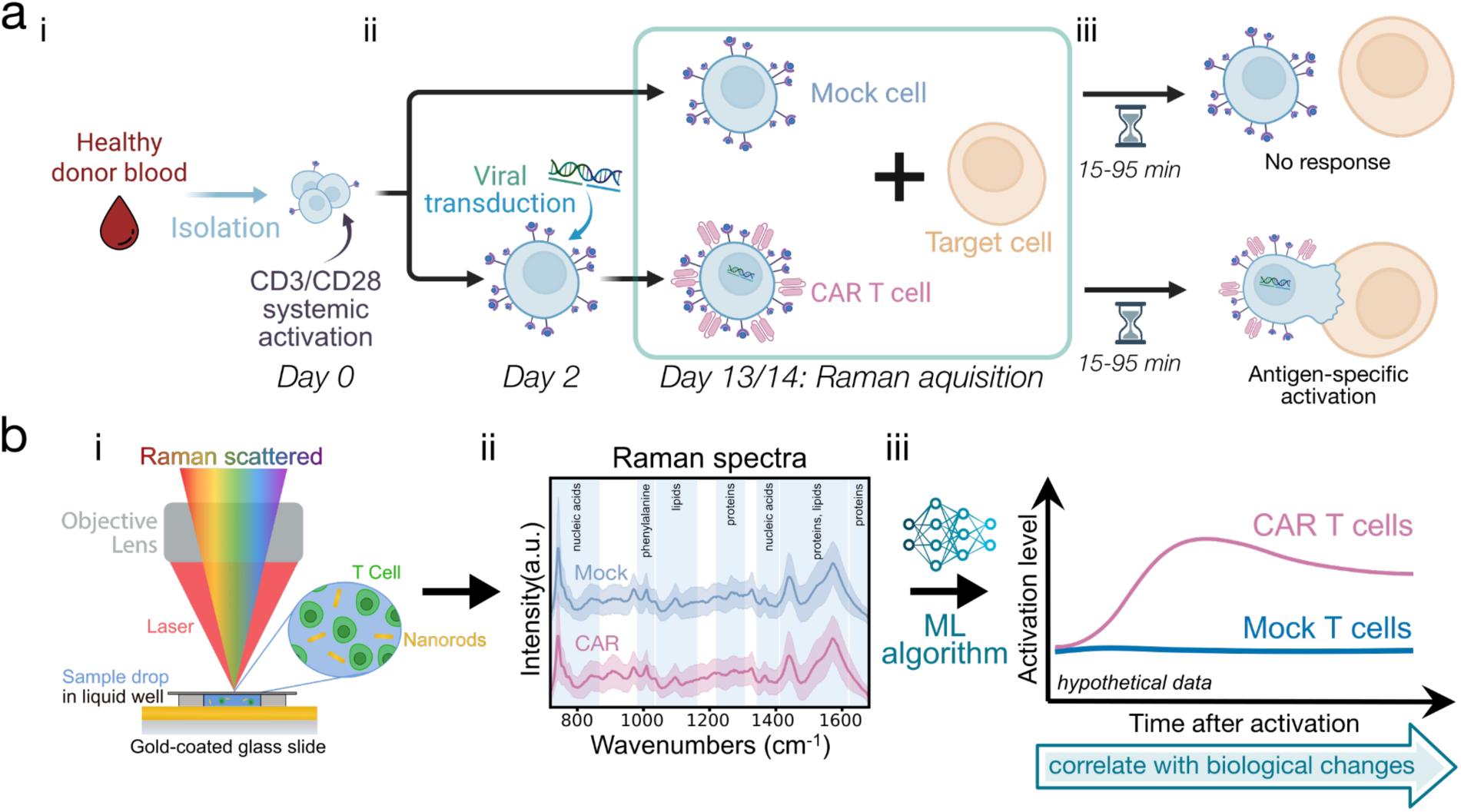
Experimental setup and workflow. **(a)** (i) T cells are isolated from healthy donor blood and activated with CD3/CD28. (ii) They are then separated into “Mock” T cells (further expanded without any introduction to viral vectors) and Chimeric Antigen Receptor (CAR) T cells, which undergo viral transduction to express CAR proteins. (iii) These two cell groups are later co-cultured with target cells expressing the protein that the CAR targets. After an incubation period, an antigen-specific activation response can be observed in the sample with CAR T cells. **(b)** To observe and analyze the cells and processes outlined in (a), we develop a Raman spectroscopy-based platform. (i) Single-cell Raman spectra are collected on a liquid well containing both gold nanorods and live cells. These spectra (ii) are used as input in a machine learning algorithm (iii), where the trained model will distinguish between CAR and Mock cells. All experiments were performed on cells day 13 or 14 after CD3/CD28 activation, and antigen-specific activation Raman measurements were collected 15-95 minutes after initiating co-culture.

## Results

### Nanorods enable rapid live-cell SERS

We generated CAR T cells from healthy donor T cells using standard activation and retroviral transduction protocols (Fig. 1a). T cells were transduced with vectors encoding either CD19.28ζ (generating CD19-CAR T cells) or GD2.4-1BBζ (generating GD2-CAR T cells).^3,25^ As a negative control, donor-matched Mock T cells were generated under the same activation and expansion conditions, but without viral transduction. Scanning electron micrographs (SEMs) of fixed, dried CAR and Mock T cells showed no major morphological differences (Fig. 2a). We assessed CAR expression by flow cytometry (Fig. 2b). Transduction efficiencies varied by donor, ranging from 55-96% for CD19-CAR T cells and 35-41% for GD2-CAR T cells (Supplementary Table 1), consistent with reported clinical manufacturing ranges of ∼30-70%.^26^

**Fig. 2:**
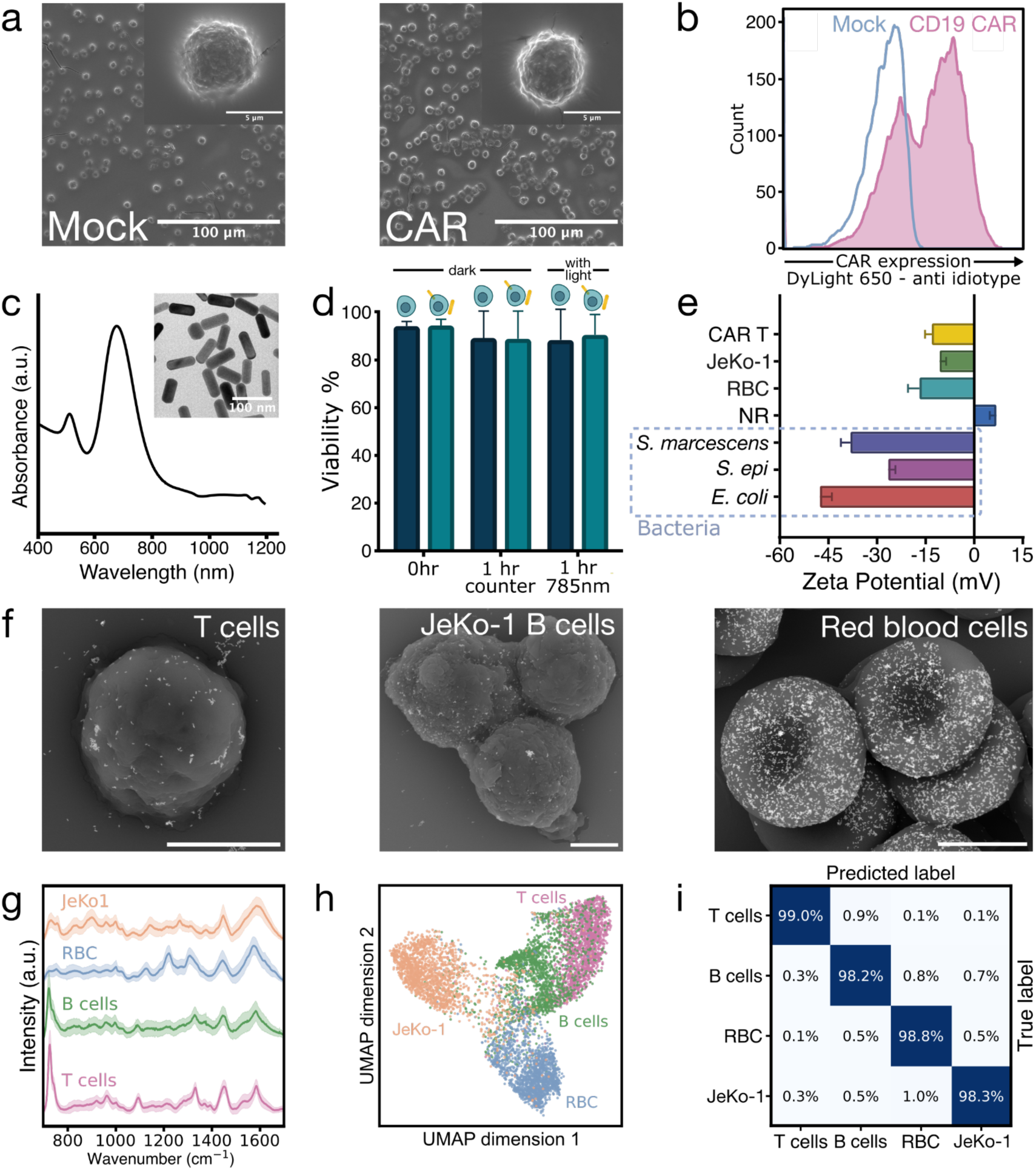
Gold nanorod-enabled surface-enhanced Raman spectroscopy (SERS) of blood cell types. **(a)** Scanning electron micrographs of Au/Pd-coated, dried Mock (left) and CAR (right) T cells. **(b)** Flow cytometry of CD19-CAR expression in healthy human donor T cells compared to Mock. **(c)** Transmission electron micrograph of gold nanorods (left) and their localized surface plasmon resonance peak at ∼670 nm (right). **(d)** Cell viability of cells incubated with and without nanorods and exposed to laser illumination for 1 hour. Conditions were performed in duplicate (n=2), with mean and standard deviation. **(e)** Surface charge (zeta potential) of blood cells, bacteria, and gold nanorods. **(f)** Scanning electron micrographs of Au/Pd-coated and dried T, JeKo-1 B, and red blood cells mixed with nanorods (lighter contrast features). Scale bars indicate 3 µm. **(g)** Mean normalized single-cell Raman spectra of live T, primary B, JeKo-1 B, and red blood cells with ±1 standard deviations (SD) (shaded). **(h)** Two-dimensional Uniform Manifold Approximation and Projection (UMAP) of Raman spectra, balanced by class. **(i)** Normalized confusion matrix displaying classification accuracy for these Raman spectra (non-reduced), generated by a Light Gradient-Boosting Machine (LightGBM) classifier with 10-fold stratified cross-validation.

Gold nanorods were selected for live-cell SERS because of their optical tunability, broad biocompatibility, and scalable synthesis. Nanorods were synthesized by seed-mediated colloidal growth to exhibit an approximate resonance peak of 680 nm, blue-shifted to minimize extinction losses in liquid under 785 nm excitation (Fig. 2c; Supplementary Fig. 1).^27^ Acridine orange/propidium iodide assays confirmed biocompatibility after 1-hour incubations with or without nanorods and with or without 785 nm illumination (Fig. 2d). A nanorod concentration of ∼385 µg/mL (nominal based on stock; Methods) ensured reproducible SERS enhancement through electrostatic association between positively-charged washed nanorods (ζ ≈ +7 mV) and negatively charged mammalian cells (ζ ≈ −14 mV on average; Fig. 2e). This approach avoided ligand-exchange steps that would introduce Raman background. The gold nanorods serve as nonspecific plasmonic enhancers that amplify intrinsic cellular vibrational signatures without targeting predefined markers. Dried SEM micrographs confirmed nanorod binding to cell surfaces (Fig. 2f), and cryogenic transmission electron microscopy verified this association under liquid conditions (Supplementary Fig. 2; Supplementary Videos 1-3).

Raman spectra were collected from live cells in a sealed liquid well chamber (Fig. 1bi). Under the same acquisition parameters, SERS spectra exhibited substantially higher intensity than non-enhanced measurements, and nanorod-only controls confirmed that dominant spectral features originated from cells (Supplementary Fig. 3). Local variations in nanorod distribution resulted in expected hotspot heterogeneity, with each spectrum capturing a snapshot of the probed cellular microenvironment. Averaging across spectra produced a representation of the overall biochemical composition.

### Human blood cell classification

As an initial demonstration of targeted-label-free live cell identification, we acquired ∼11,000 single-cell Raman spectra from four cell types — T cells (untransduced and not systemically activated), primary human B cells, mantle cell lymphoma B cell line (JeKo-1), and red blood cells (RBCs), all suspended in phosphate-buffered saline (PBS) (Fig. 2g; Supplementary Table 2; Methods). Uniform Manifold Approximation and Projection (UMAP) visualization of preprocessed spectra identified clustering by cell type (Fig. 2h; Supplementary Fig. 4).^28^ ML models trained on the full non-reduced spectral dataset achieved 98.6% classification accuracy using 10-fold stratified cross-validation (CV) (Fig. 2i; Supplementary Table 3).

Discriminating peaks correspond with known Raman biochemical features resulting from blood cell identity (Supplementary Fig. 5; Supplementary Table 4),^20,29–35^ including those from DNA/RNA (adenine ring breathing 725-730 cm^−1^, DNA backbone PO_2_^−^ stretch 1,095-1,100 cm^−1^), phenylalanine (1,004 cm^−1^), cytochrome c (750 cm^−1^), hemoproteins (1,128, 1,225, 1,565-70, 1,620-25 cm^−1^), general protein bands (1,140-1,160, 1,620-65 cm^−1^), and lipids (1,330, 1,460 cm^−1^). The adenine-associated peak at ∼725 cm^−1^ had the largest contribution to classifying T cells, primary B cells, JeKo-1 B cells, and red blood cells (Supplementary Fig. 5). Its reduced presence in both B cell populations relative to T cells is consistent with a smaller nuclear contribution within the sampling volume, potentially influenced by differences in cell size and nucleus-to-cytoplasm geometry. Mature erythrocytes, lacking nuclei and mitochondria, show minimal nucleic acid and cytochrome c signatures but strong hemoprotein bands. Any residual intensity around 724 cm^−1^ in the RBC spectra likely arises from hypoxanthine, which previous SERS studies found accumulates during blood storage.^32^ Relative to T cell spectra, JeKo-1 B cell spectra exhibit more intense peaks associated with phenylalanine and other general proteins. This may partly reflect their larger size, as intensity can reflect protein concentration. Additionally, transformed cells such as JeKo-1 B cells are in a proliferative state, which may contribute to increased protein production.^30^ Overall, primary B cell spectra tend to fall between T and JeKo-1 cells across multiple discriminative features.

### Distinguishing CD19-CAR from Mock T cells

Having demonstrated blood cell type discrimination, we next asked whether SERS could distinguish more subtle molecular differences arising from T cell engineering. We acquired ∼37,000 single-cell spectra from live anti-CD19-CD28-CD3ζ CAR (CD19-CAR; 55-96% transduction efficiency) and donor-matched non-engineered Mock T cells across nine healthy donors (Fig. 3a, top; Methods). Training models on each donor individually achieved an average donor-level 10-fold CV classification accuracy of 75.0% (Supplementary Fig. 6). Analysis of intensity differences (Fig. 3a, bottom) and feature importances (Fig. 3b) revealed that CD19-CAR T cell spectra exhibited consistent enrichment in protein and aromatic bands (880-900, 1,020-1,030, broadly 1,560-1,610 cm^−1^),^29,30,33,35^ whereas Mock cells showed stronger nucleic acid signatures (725-735, 1,090-1,100, 1,330-1,340 cm^−1^).^20,29–31,33,36^ Additionally, we observed increased emphasis at 717-721 and 870-880 cm^−1^ for CD19-CAR T cells, consistent with membrane phosphatidylcholine (phospholipid) headgroup modes.^37^ Consistent with these protein and aromatic signature differences, single-cell RNA sequencing of CD19 CAR T cell infusion products from seven patients showed enrichment in CAR+ cells for proteostasis-associated pathways, including protein folding, as well as cell cycle processes (Supplementary Fig. 7c-d).

**Fig. 3:**
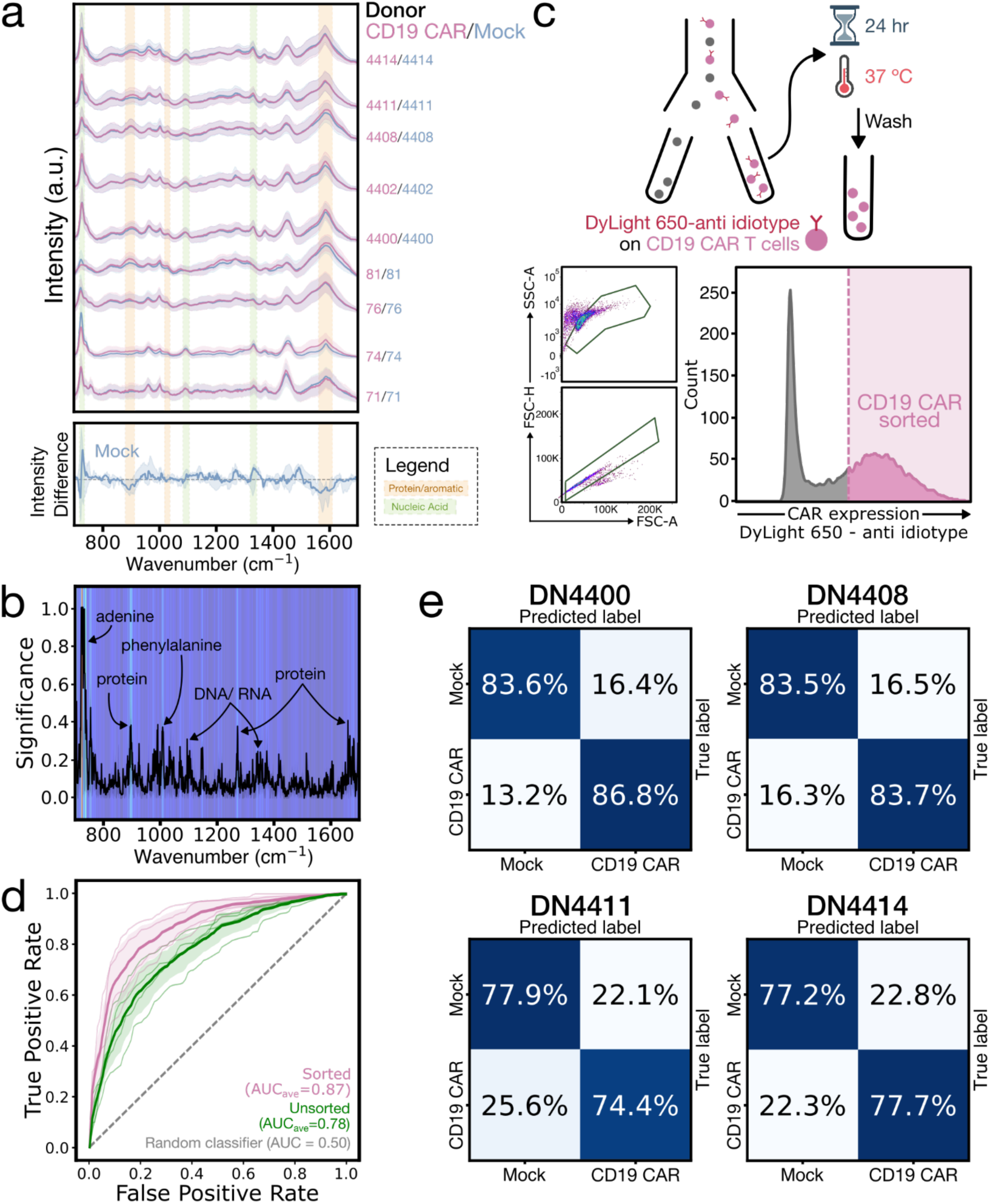
CD19-CAR and Mock T cell SERS classification. **(a)** Waterfall plot of mean normalized SERS spectra of live single CD19-CAR (pink) and Mock (blue) T cells with ±1 SD (shaded), separated by donor (identifier labeled on the right) (top). Median donor-normalized spectral difference of Mock cells relative to CAR with 25-75% quantile range (bottom). The grey dashed line marks zero difference. Colored spectral bands highlight CAR-enriched protein/aromatic (gold) and Mock-enriched nucleic acid (green) regions. Over 2,000 spectra were acquired per donor. **(b)** Mean feature importance plot depicting the contribution of each spectral dimension (wavenumber) to classification, both through the curve and the heatmap background. Significant peaks are labeled with their biochemical vibrational modes. **(c)** Cell sorting schematic (top). CD19-CAR T cells were labeled with DyLight 650-conjugated FMC63 anti-idiotype and flow sorted to achieve a 100% CAR positive sample. After overnight incubation and washing, the labels were effectively removed and cells were analyzed by Raman spectroscopy. Flow cytometry plots show gating for single cells (bottom left) and DyLight 650 labeling intensity (bottom right). **(d)** Receiver operating characteristic (ROC) curves comparing classification performance of unsorted (green) and sorted (pink) CAR T cells versus Mock. Thick lines show the mean across four donors with ±1 SD and thin lines show individual donor curves. The grey dashed line represents a random classifier. Areas under the curve (AUC) are given in the legend. **(e)** Normalized confusion matrices for 4 of 9 donors displaying the classification accuracy for sorted CD19-CAR and Mock Raman spectra using a LightGBM classifier with 10-fold stratified cross-validation.

Together, this suggests that CAR expression shifts the probed cellular volume towards membrane, protein, and aromatic contributions, while Mock spectra show a greater dominance of nucleic acid bands. The increased protein and aromatic prominence in CD19-CAR T cells may reflect contributions from the CAR protein itself, which contains aromatic amino acids (tryptophan, tyrosine, and phenylalanine) in its scFv domain.^38^ Additionally, previous studies suggest that CAR proteins can self-assemble into raft-like domains stabilized by microvilli, which may locally concentrate membrane protein and phospholipid signatures, even in non-target-engaged cells.^39^ Conversely, Mock cells may produce spectra with relatively greater nuclear volume sampling, consistent with a more quiescent state and a less protein-rich membrane.

These trends were further validated using Raman spectra acquired without gold nanorods (∼7,800 spectra, three donors, 25-second acquisitions; Methods), which remove surface-enhancement and substrate-specific biases. As expected, the adenine-associated bands were less pronounced in the absence of preferential gold nanorod binding, but the key class-dependent trends were preserved (Supplementary Fig. 8),^40^ supporting that these are genuine biochemical differences rather than SERS sampling bias.

Classification accuracy was further improved by isolating CD19-CAR positive cells from the ∼4-45% untransduced fraction observed across all nine donors (reflecting typical lentiviral transduction efficiency) via fluorescence-activated cell sorting (FACS), followed by label-clearing step to minimize contributions from fluorescent antibody tags (Supplementary Fig. 9). Using models trained on the purified CAR T cells (∼10,200 spectra across four donors), average donor-level classification accuracy increased from 70.3% to 80.6%, with all donors showing clear improvements in accuracy and area under the receiver operating characteristic curve (AUC) (Fig. 3d-e). This further demonstrates the detection sensitivity of this approach to CAR expression frequency.

### Time-resolved SERS of antigen-specific activation

We next investigated whether our SERS-based method could capture dynamic functional changes during antigen-specific activation. CD19-CAR and unstimulated (Unstim; isolated donor T cells without CD3/CD28 activation) T cells were co-cultured with CD19+ JeKo-1 B cells to induce antigen-specific activation of CAR T cells (Fig. 4a). Unstimulated T cells provided a baseline control for time-dependent spectral drift, ensuring we could isolate effects arising solely from activation and CAR expression. Cells were co-incubated at a 3:1 effector-to-target ratio that enriches for activation events.^41^ Spectra were acquired semi-continuously to monitor population-level activation dynamics (Fig. 4b; Methods). UMAP visualization of ∼1,800 Raman spectra per donor across two donors revealed distinct clustering between CD19-CAR and Unstim conditions (Fig. 4c), with ML models achieving an average 94.6% classification accuracy via 10-fold stratified cross-validation (Fig. 4d).

**Fig. 4:**
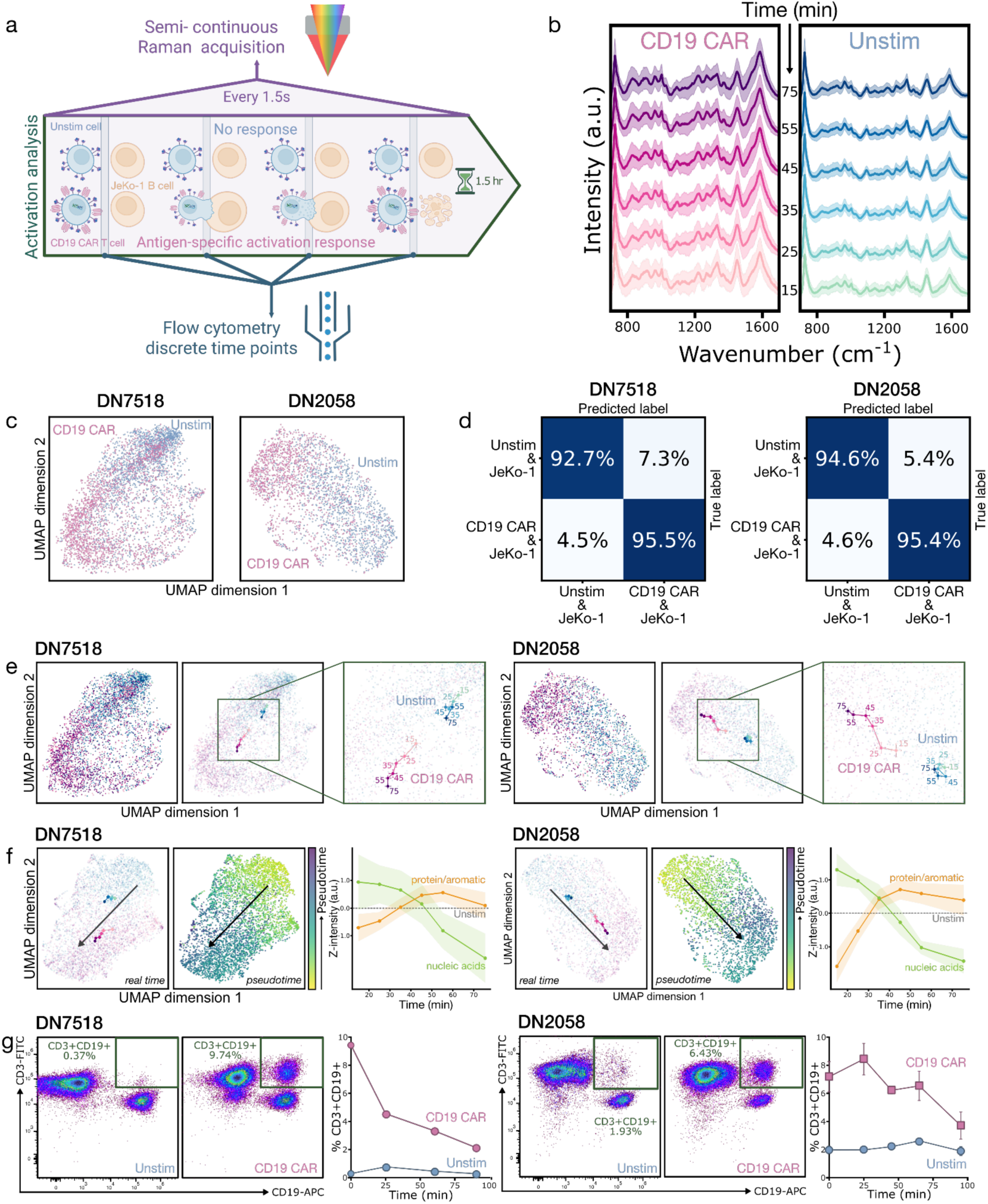
Antigen-specific activation analysis of CD19-CAR T cells by time-resolved SERS. **(a)** Diagram of experimental workflow. CD19-CAR and unstimulated T cells were co-cultured with CD19+ JeKo-1 B cells, where only CAR T cells undergo antigen-specific activation. Semi-continuous Raman acquisition was paired with discrete flow cytometry time points. **(b)** Mean normalized live-cell SERS spectra of CD19-CAR (pink) and unstimulated (blue) T cells co-cultured with JeKo-1 B cells, averaged in 10-20 minute segments (labels indicate segment start time from 15-75 minutes) over two donors. **(c)** Two-dimensional UMAPs of the Raman spectra plotted in (b). **(d)** Normalized confusion matrices displaying classification accuracy combined across the 1.5-hour time course for two donors. Accuracies were generated using a LightGBM classifier with 10-fold stratified cross-validation. **(e)** UMAPs of Raman spectra from two donors. Points are labeled by CAR (pink) and unstimulated (blue), with color shading from light to dark indicating time from 15 to 75 minutes (left). The centroids of each time segment are on the UMAP projection (middle). When zoomed in and labeled with their starting time point, they show a trajectory during activation (or lack thereof for unstimulated T cells) (right). **(f)** Pseudotime trajectory analysis of CAR T cell activation for two donors. UMAPs with real-time centroid trajectories, where darker hues indicate later time points, as in (e) (left). The pseudotime trajectories are visualized on UMAPs by a pseudotime color scale (yellow to purple) (middle). The arrows indicate the overall temporal direction. Co-cultured CAR T cell Raman peak intensity trends over real time for nucleic acid (green) and protein/aromatic (orange) bands, normalized to unstimulated cells (right). Curves represent median trajectories with shaded ribbons indicating 95% confidence intervals (n = 331–1,196 spectra/bin). **(g)** Flow cytometry validation of activation, measured by the CD3+CD19+ population (in the green box) for two donors. The flow fluorescence plots show representative distributions at the initial time point (marked as minute 0 on the right plot) (left, middle), and percentages of CD3+CD19+ cells are plotted over four discrete time points (right).

Population-level comparisons aggregated across time (∼7,400 spectra total) displayed differences reflecting both CD19-CAR expression and activation (Supplementary Fig. 10). Target-engaged CD19-CAR T cell spectra showed greater protein and aromatic prominence at 1,002 (phenylalanine), 1,031, and broadly 1,500-1,660 (amide I and II) cm^−1,29,30,32,33,35^ as well as increased signal in the 1220-1280 cm^−1^ (amide III) region. In contrast, Unstim T cells, in an even more quiescent state than Mock, had relatively enriched nucleic acid-associated signatures at 726 (adenine), 964, 1,095 (DNA backbone), 1,330-40, and 1,453 cm^−1^.^20,29–31,33,36^

These trends align with the CD19-CAR versus Mock differences we observed in individual cells — where SERS of protein-rich CAR T cells showed comparatively reduced nuclear sampling — and are also consistent with prior non-SERS Raman studies of T cell activation. The relative decrease in nucleic acid peaks during activation has been theorized to arise from chromatin decondensation, leading to a more open euchromatic architecture and therefore a lower local DNA concentration within the Raman probe volume.^20,21,29^ The broad increase in protein-related signatures for target-engaged CD19-CAR T cells is consistent with changes in protein composition and organization reported during T cell activation. During antigen-specific activation, T cells undergo synapse formation involving increased translation and protein synthesis, receptor recruitment to the membrane, cytoskeletal polarization, and membrane reorganization (including receptor aggregation).^29,39^ The co-cultured CD19-CAR T cell spectra showed increased relative signal within the 1220-1280 cm^−1^ region corresponding to amide III bands that reflect protein secondary structure and local environment and are sensitive to structural order.^33,35^ Increases in this band may indicate changes in protein backbone structure and bonding environment. Prior visible-excitation Raman studies have reported carotenoid-associated bands distinguishing T cells from B cells and decreasing with T cell activation.^20,31,42^ In our near-infrared (IR) SERS measurements, we did not observe prominent carotenoid contributions, consistent with much lower carotenoid Raman cross-sections in the near-IR^43^ and with potential additional effects from SERS sampling geometry.

Spectra from T- and JeKo-1 B-cell co-cultures captured the different stages of T-cell activation and B-cell cell death. To visualize activation progression, mean UMAP positions (“centroids”) for each condition were plotted over time (Fig. 4e). Although UMAP does not preserve global distances, qualitative shifts in the embedding can illustrate cluster evolution. For both donors, CD19-CAR centroids showed clear progression with time, indicating biochemical changes during activation, while Unstim centroids appeared comparatively stationary. As a complementary analysis, we applied an unsupervised pseudotime method that arranges spectra into a continuous trajectory based on their biochemical similarity (Fig. 4f, middle; Methods). This produced a pseudotime progression that strongly correlated with true experimental time (Spearman’s rank correlation of 1.00 and 0.94 for two donors; Fig. 4f, left; Supplementary Fig. 11).^44^ Notably, this method recovered an activation trajectory without using time labels, demonstrating that the Raman spectra inherently contain the temporal structure of activation.

Temporal tracking of spectral bands for both donors (Fig. 4f, right) displayed an initial rise in protein and aromatic bands during the first hour, followed by a partial decline (Spearman’s rank correlation of 0.83 and 0.6). Nucleic acid bands decreased monotonically, with a slight leveling after an hour (Spearman’s rank correlation of −1.00). The early trends are consistent with known activation processes mentioned above.^29,39^ Later leveling likely reflects a transition towards a steady effector state or population-level averaging over activated, interacting, and dying T and JeKo-1 B cells. We benchmark our findings against an established activation readout by performing parallel flow cytometry measurements of CD3+ CD19+ events — indicative of multi-cell events with interacting T and JeKo-1 B cells or trogocytosis.^38^ Activated CD19-CAR co-cultures showed a high early population of events that declined over time, while Unstim controls started low and remained unchanged (Fig. 4g; Supplementary Fig. 12). These measurements complement the Raman-derived trajectories, notably the later-time leveling, and provide an orthogonal validation of the activation dynamics captured with our targeted-label-free approach. Note that this flow assay detects conjugates, which may resolve more quickly, rather than underlying biochemical changes. Overall, these findings demonstrate that our approach enables dynamic live cell phenotyping with temporal resolution limited only by Raman acquisition time, in contrast to conventional flow cytometry, which captures fewer, more discrete time points and requires perturbative labeling and time-intensive processing.^21^

### Generalizability to a tonic signaling CAR (GD2-CAR)

To validate the robustness of this methodology across different CAR architectures, we generated and analyzed anti-GD2-4-1BB-CD3ζ CAR (GD2-CAR) T cells using the same activation, transduction, and expansion protocol as for CD19-CAR T cells. Donor-matched Mock T cells were generated in parallel and, as before, were activated and expanded but left untransduced. GD2-CAR T cells differ from CD19-CAR T cells in receptor antigen recognition and costimulatory domains (Fig. 5a). Additionally, GD2-CAR T cells exhibit high levels of tonic signaling, where clustering of the CAR surface proteins triggers basal ζ-chain phosphorylation and downstream metabolic activation independent of antigen recognition.^45^

**Fig. 5:**
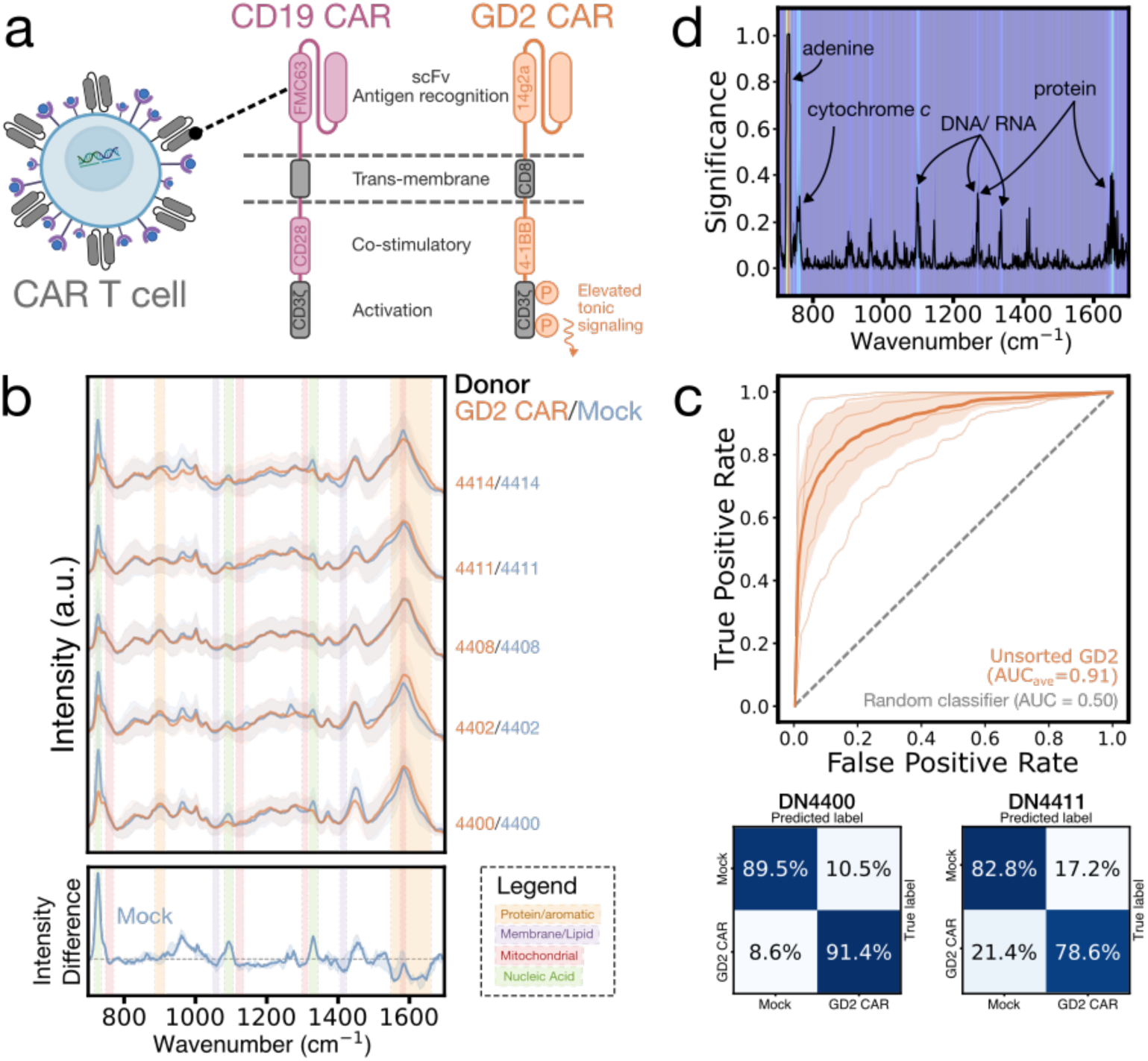
GD2-CAR and Mock T cell SERS classification. **(a)** Diagram comparing CD19-CAR (pink) and GD2-CAR (orange). Differences in antigen recognition and co-stimulatory domains are highlighted, with GD2-CAR showing elevated tonic signaling due to ζ-chain phosphorylation. **(b)** Waterfall plot of mean normalized SERS spectra of live single GD2-CAR (orange) and Mock (blue) T cells with ±1 SD (shaded), separated by donor (identifier labeled on the right) (top). Median donor-normalized spectral difference of Mock cells relative to CAR with 25-75% quantile range (bottom). The grey dashed line marks zero difference. Colored spectral bands highlight CAR-enriched protein/aromatic (gold), membrane/lipid (purple), and mitochondrial (red) regions, and Mock-enriched nucleic acid regions (green). **(c)** ROC curves displaying classification performance of GD2-CAR T cells versus Mock (top). Thick lines show the mean across four donors with ±1 SD and thin lines show individual donor curves. The grey dashed line represents a random classifier (AUC = 0.5). AUCs are given in the legend. Representative normalized confusion matrices for 2 of 5 donors displaying classification accuracy of Raman spectra using a LightGBM classifier with 10-fold stratified cross-validation (bottom). **(d)** Mean feature importance plot depicting the contribution of each spectral dimension (wavenumber) to classification, both through the curve and the heatmap background. Significant peaks are labeled with their biochemical vibrational modes.

Across five donors (∼11,800 total spectra, Fig. 5b; Methods), donor-level classification accuracies averaged 85%, with average AUC improving from 0.78 (unsorted CD19-CAR) to 0.91 (Fig. 5c; Supplementary Fig. 13). Spectral analysis (Fig. 5b, bottom; Fig. 5d) showed that GD2-CAR T cells, like CD19-CAR T cells, displayed enriched protein and aromatic bands (892 and broadly around 1,600 cm^−1^)^30,33,35^ and weaker nucleic acid bands (728, 1,094, 1,335 cm^−1^)^20,29–31,33,36^. Additionally, GD2-CAR T cells have elevated mitochondrial signatures at 755, 1,130, 1,310, and 1,584 cm^−1^, consistent with cytochrome c-associated mitochondrial expansion.^20^ Orthogonal representative measurements — mitochondrial mass flow cytometry and mitochondrial fraction protein quantification — show higher mitochondrial content in GD2-CAR+ cells (Supplementary Fig. 14). These trends align with expected tonic signaling driven increases in protein synthesis, cellular stress, and mitochondrial content. Additionally, the 4-1BB costimulatory domain in the GD2-CAR variant favors fatty acid oxidation and mitochondrial biogenesis, resulting in elevated levels of cytochrome c complexes.^46^ Conversely, CD28 costimulation (found in the previous CD19-CAR construct) biases cells towards aerobic glycolysis (Warburg metabolism), which bypasses mitochondrial oxidation for energy production and leads to minimal mitochondrial expansion.^46,47^ This explains the lack of strong mitochondrial significance when comparing CD19-CAR and Mock T cell spectra. The higher classification accuracy for GD2-CAR demonstrates that as phenotypic changes increase, spectral separability strengthens. These results demonstrate that SERS can distinguish engineered T cell states driven by differences in a single membrane protein type, and can do so across distinct CAR architectures. This emphasizes the potential of our approach for profiling diverse engineered cell therapies.

### Longitudinal SERS profiling of CD19-CAR T therapy patient samples

To explore the feasibility of our SERS-ML workflow in clinical samples, we applied it to a four-patient cohort of longitudinal CD19-CAR T therapy samples collected pre-infusion and at days 7 and 21 post-infusion, referred to as Pre, Peak, and Late, respectively, where day 7 corresponds to the expected peak CAR T cell expansion window and day 21 to the late expansion timepoint.^48^ Samples were analyzed as mixed PBMCs at all timepoints. Post-infusion samples were additionally separated using FACS into three populations for Raman analysis: CD3+CAR+, CD3+CAR-, and Dump+, where the Dump+ population consisted of cells captured by a lineage channel containing CD19, CD14, and CD235a antibodies marking B cell, monocyte, and red blood cell populations, respectively. The experimental workflow is shown in Fig. 6a. In addition to single-cell SERS profiling, we analyzed matched cytometry by time of flight (CyTOF) data from a prior study,^48^ as well as CAR T cell frequencies acquired during FACS, to provide orthogonal context for patient and timepoint-specific immune state (Supplementary Fig. 15; Supplementary Table 5). Overall, the Raman-analyzed patient samples spanned multiple heterogeneous and therapeutically relevant immune states, motivating patient-specific Raman analyses.

**Fig. 6:**
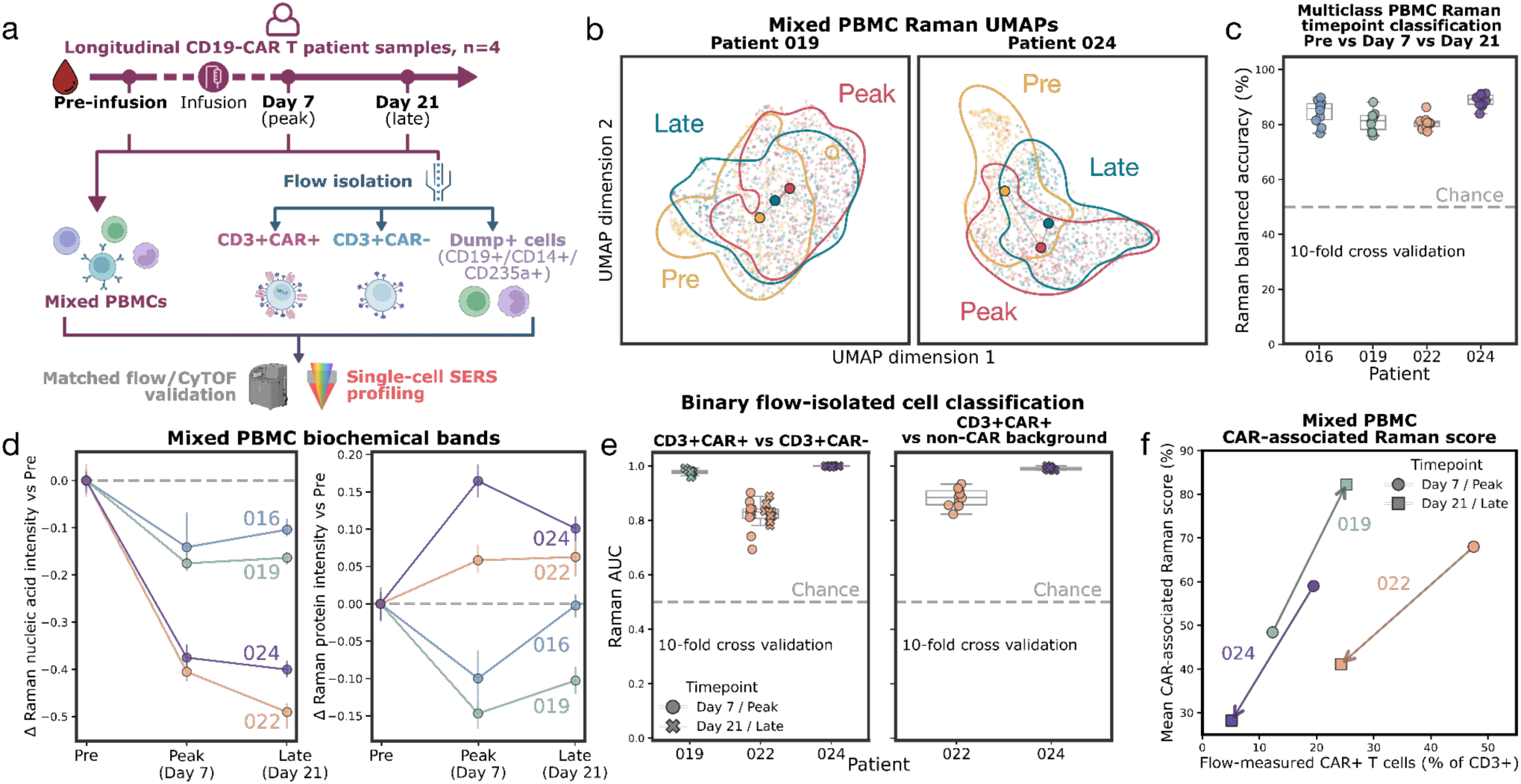
Longitudinal CD19-CAR T patient sample classification. **(a)** Experimental workflow. Longitudinal PBMC samples were collected from four CD19-CAR T therapy-treated patients before infusion (Pre) and after infusion at Day 7 (Peak) and Day 21 (Late). Mixed PBMCs were analyzed at all timepoints, and post-infusion samples were additionally FACS sorted into CD3+CAR+, CD3+CAR-, and Dump+ (CD19, CD14, and CD235a) populations. Flow-isolated and mixed PBMC samples were analyzed by single-cell SERS and interpreted alongside matched flow cytometry and CyTOF data.^48^ **(b)** Representative UMAPs of mixed PBMC Raman spectra colored by timepoint (full cohort UMAPs are shown in Supplementary Fig. 16). Faded points represent individual spectra, larger markers indicate timepoint medoids, and contours highlight higher-density regions for each timepoint. **(c)** Multiclass classification performance for mixed PBMC Raman spectra by timepoint for each patient. Models classified spectra as Pre, Day 7/Peak, or Day 21/Late. Each point represents the balanced accuracy from one CV fold (*n* = 10 folds), with distributions summarized by box plots. The grey dashed line indicates chance. Confusion matrices summarizing the performance of each model can be found in Supplementary Figure 16. **(d)** Longitudinal nucleic acid-associated (left) and protein-associated (right) Raman band intensities in mixed PBMCs, shown relative to each patient’s matched Pre sample (patients differentiated by color). Points show medians; error bars indicate 95% bootstrap confidence intervals. **(e)** Binary classification performance for flow-isolated Raman spectra. Dot plots show CD3+CAR+ versus CD3+CAR-T cells (left) and CD3+CAR+ versus non-CAR background cells (right). Each point represents AUC from one CV fold (*n* = 10 folds). Models were trained separately for each patient/timepoint grouping with sufficient spectral counts. Peak and Late classifications are marked by circles and crosses, respectively. **(f)** CAR-associated Raman score analysis in mixed PBMCs. Patient-specific classifiers were trained to distinguish Pre mixed PBMC Raman spectra from post-infusion flow-isolated CD3+CAR+ spectra with sufficient spectral counts, then applied to Peak and Late mixed PBMC spectra. Mean classifier output probabilities, defined as the CAR-associated Raman score, are plotted against flow-measured CAR+ T cell frequency within CD3+ cells. Arrows indicate Peak-to-Late direction.

Across four patients and three timepoints, ∼8,200 SERS spectra were acquired from mixed PBMC samples (Supplementary Table 6). Patient-specific ML models classified Pre, Peak, and Late spectra with an average accuracy of ∼84%. These timepoint-associated spectral differences likely reflect the combined effects of lymphodepletion, lymphocyte proliferation, CAR T cell expansion, cellular activation, and broader post-infusion immune remodeling. Representative UMAPs of PBMC spectra showed patient-specific longitudinal structure (Fig. 6b). Fig. 6c shows 10-fold cross-validated classification performance for predicting sample timepoint for each patient. Full spectra, UMAP embeddings, and confusion matrices are provided in Supplementary Fig. 16. Unlike controlled healthy donor CAR versus Mock T cell experiments, longitudinal PBMC samples from CAR T therapy patients vary simultaneously in CAR T cell frequencies, cell-type composition, treatment history, and multiple functional state phenotypes. Qualitatively, the Pre timepoint clusters often formed the most distinct population in UMAP embeddings, while the Late timepoint tended to show broader redistribution, consistent with heterogeneous post-infusion immune remodeling and partial overlap between later and baseline-like cell states.^49^

We next tracked the timepoint-specific biochemical trends in nucleic acid-associated bands (728, 1,094, 1,335 cm^−1^)^20,29–31,33,36^ and protein and aromatic bands (880-900, 1,002, 1,020-1,030, 1,220-1,290, 1,560-1,610, and 1,630-1,660 cm^−1^),^29,30,33,35^ (Fig. 6d). Across patients, post-infusion nucleic acid-associated Raman intensity tended to decrease relative to each patient’s matched Pre baseline, whereas protein/aromatic-associated trends were more patient-dependent. Contrasts between nucleic acid- and protein/aromatic-associated regions, single-spectrum distributions, and selected additional biochemical band trends are provided in Supplementary Fig. 17. These findings support the interpretation that longitudinal Raman spectra capture integrated immune state changes during the therapeutic lifecycle rather than a single CAR-specific molecular feature or isolated marker change.

In addition to mixed PBMC spectra, we acquired Raman data from flow-isolated CD3+CAR+, CD3+CAR-, and Dump+ populations. Analyzing only patient/timepoint groupings with sufficient spectral counts (Supplementary Table 6; Methods), CD3+CAR+ spectra were distinguishable from CD3+CAR-T cells with an average grouping-specific accuracy of 86% and an average AUC of 0.93 (Fig. 6e; Supplementary Fig. 18). As a separate classification task, CD3+CAR+ spectra were compared against a broader non-CAR background class that pooled CD3+CAR-T cells with Dump+ non-T cells. CD3+CAR+ cells were also distinguishable from this pooled background with an average accuracy of 88% and an average AUC of 0.95. These results extend the engineered versus endogenous classification demonstrated in donor-derived models to clinically relevant patient-derived samples.

Finally, we asked whether ML models could identify CAR-associated spectral signatures within mixed PBMC samples. Patient-level classifiers were trained to distinguish matched pre-infusion mixed PBMC spectra from flow-isolated CD3+CAR+ spectra collected after infusion, and then were applied to Peak and Late mixed PBMC spectra. For this analysis, post-infusion CD3+CAR+ reference spectra were included only from patient/timepoint combinations with sufficient spectral counts (Methods). The resulting predicted probability was defined as a CAR-associated Raman score and compared with matched thawed flow measurements from the same Raman-analyzed samples. This score was interpreted as a measure of CAR-associated spectral similarity rather than a calibrated estimate of CAR+ T cell frequency (Fig. 6f; Supplementary Fig. 19). CAR-associated Raman score changes from Peak to Late timepoints were directionally concordant with matched flow-measured CAR+ T cell frequency changes, supporting that the classifier detected CAR-associated spectral structure within mixed PBMC samples.

## Discussion

We developed a SERS- and ML-based method for semi-continuous, non-destructive identification and functional-state monitoring of live CAR T cells without molecularly targeted labels. Using a plasmonic platform, we collected rich molecular spectra that capture biochemical and biophysical changes resulting from blood cell type, engineered receptor expression, and activation processes. We demonstrated these capabilities in donor-derived models and extended the workflow to longitudinal CAR T therapy patient samples. Taken together, these findings demonstrate the value of this technique for probing cell identity and multidimensional functional-state information, including dynamic processes with temporal specificity. In this work, we used a conservative 1.5-second acquisition time to ensure consistent high signal-to-noise across experimental conditions. This parameter can be readily reduced to the sub-second regime with speed-optimized configurations such as higher excitation power, greater plasmonic/nanophotonic enhancement, and improved optical throughput.^13,17^

Our spectral interpretations link vibrational molecular signatures to known CAR T cell biology, demonstrating the strength of Raman as a biochemistry-based readout. However, several challenges remain before achieving translational impact. Donor variability, transduction heterogeneity, and limited sample size currently hinder model generalizability across unseen donors and patients. This limitation does not preclude longitudinal single-patient monitoring, where patient-specific or baseline-referenced models are sufficient. The patient-derived experiments provide an initial extension of this approach into clinically heterogeneous samples, which introduce additional complexity from prior treatment history, cell-type composition, and immune state phenotypes.^48^ The CAR-associated Raman score should therefore be interpreted as a classifier-derived measure of spectral similarity rather than calibrated quantification of CAR+ T cell frequency. Broader donor-agnostic generalization is valuable for applications like manufacturing quality control and multi-site deployment, where training patient-specific models may be impractical. Improving generalizability necessitates spectral libraries spanning multiple donors, patient cohorts, receptor constructs, and cell functional states. While SERS sampling bias and hotspotting can contribute to spectral variability, we have shown that complementary non-SERS measurements can help validate underlying biochemical features. Alternative SERS platforms, such as nanofabricated metasurfaces, have the potential to reduce sampling variability, improve reproducibility, and completely omit any cell-surface nanostructures. Our co-culture measurements inherently averaged across interacting and dying T and JeKo-1 B cells, while PBMC measurements averaged across a highly heterogeneous mixture of cell types. Further studies tracking a single cell-cell interaction, either in microfluidics or an isolation well, would help reduce fluctuations resulting from variations in cell state and type. Additional parallel biological assays could help deepen our spectral band interpretations (*e.g.,* information on metabolites, cytokines, or cell architecture).

A concrete translational next step would be a larger prospective clinical validation study pairing Raman profiling with standardized immune monitoring. This study should include pre-infusion CAR T products and longitudinal patient PBMC samples from a larger, clinically diverse cohort, with matched flow cytometry, single-cell sequencing, cytokine assays, toxicity grading, and treatment-response outcomes. This experiment would calibrate Raman-associated signatures against orthogonal immune measurements, test model generalization, and determine whether early Raman measurements or dynamic spectral trajectories provide actionable information about therapeutic efficacy and toxicity.

Our approach’s rapid acquisition rate and minimal sample preparation make it well suited for future applications in quantifying proliferation kinetics, studying CAR-related toxicity, informing therapeutic decisions, and screening cells for functional phenotypes such as activation and, potentially, exhaustion. Our longitudinal patient sample results provide an initial step towards these translational applications by demonstrating that the same SERS-ML platform can span key areas of the therapeutic lifecycle, from engineered cell characterization relevant to manufacturing quality control to longitudinal PBMC analysis relevant to post-infusion monitoring. Engineered cells have transformative potential not only in cancer immunotherapy but also in autoimmune disease treatment. Both antitumor and immune-tolerizing engineered cell therapies (e.g., CAR T regulatory cells) would benefit from a unified approach to real-time monitoring of cell persistence, suppressive capacity, and functional stability. Beyond CD19- and GD2-CARs, the CAR protein makeup is subject to a large amount of variation. Future work can extend this approach across receptor constructs, including differences in intracellular costimulatory or signaling domains (e.g., CD3ε, CD4, CD8α) and extracellular CAR targets such as BCMA-CAR and CD22-CAR.^1^ Integrating this approach with microfluidic sample digitization could help create an automated, high-throughput point-of-care device for single-cell Raman profiling of engineered cells in blood.

To our knowledge, this work represents the first demonstration of Raman spectroscopy for single-cell phenotyping and dynamic functional monitoring of CAR T cells, extending from donor-derived models to longitudinal clinical samples. Our approach is a novel, cross-disciplinary convergence of microscopy, nanophotonics, ML, biochemistry, and immunotherapy. As a rapid, non-destructive, and high-information-content technique, this method provides a foundation for gaining new insight into engineered cell mechanisms and helping ensure the development of safer and more effective CAR T cell therapies.

## Methods

### Mammalian cell culture

The JeKo-1 human mantle cell lymphoma line (ATCC) was obtained as a gift from the Mackall Lab (Stanford, CA). Cells were cultured in complete growth medium — RPMI 1640 Medium with GlutaMAX supplement (Thermo Fisher Scientific), 10% heat-inactivated fetal bovine serum (FBS; R&D Systems), and 1% penicillin-streptomycin (10,000 U/mL; Gibco) — at 37°C, supplied with 5% CO2. Media was changed every two or three days. De-identified human red blood cells (K_2_ EDTA anticoagulant) were acquired from BioIVT and were stored at 4°C until use. De-identified primary human CD19+ CD20+ B cells were obtained cryopreserved from BioIVT and thawed on the day of measurement.

### CAR T cell generation and culture

De-identified human T cells were acquired through our institution’s blood bank and retrovirally transduced with CD19-CAR^25^ or GD2-CAR^3^ constructs as previously described. In brief, T cells were activated with the TransACT anti-CD3/anti-CD28 reagent (Miltenyi Biotech) and 100IU/mL rhIL-2 (PeproTech) for 48 h before retroviral transduction. On day 6 or 7, transduction efficiency was assessed via flow cytometry. Cells were labeled with huCD3 (BUV395; BD; 563546), huCD4 (BUV737; BD; 612748), huCD8 (Spark Blue 550; Biolegend; 344760), LIVE/DEAD fixable viability dye (Near IR; Invitrogen; L10119), and the respective CAR anti-idiotype conjugated to DyLight650 (as per manufacturer’s protocol; Invitrogen). Cells were cultured in complete culture medium supplemented with 10 ng/mL rhIL-2 at 37°C with 5% CO_2_, with media changes every two or three days.

### Clinical patient samples

Cryopreserved PBMC samples were obtained from four patients treated with standard-of-care axicabtagene ciloleucel CD19-CAR T cell therapy for large B cell lymphoma. This study analyzed patients with cohort identifiers 016, 019, 022, and 024 from the prior study^48^. All patient samples were collected under an institutional review board (IRB)-approved protocol with patient-informed consent for research use, and samples were de-identified before analysis (IRB no. 43375). Longitudinal samples were collected pre-infusion (“Pre”), day 7 post-infusion (“Peak”), and day 21 post-infusion (“Late”). The Pre samples were collected before lymphodepletion, approximately 5 days before CAR T cell infusion. The Peak label denotes the expected time of peak CAR T cell expansion rather than the measured maximum CAR+ T cell frequency for each patient. Cryopreserved PBMCs were thawed and analyzed the same day by both Raman spectroscopy and FACS.

### Nanorod synthesis and characterization

Gold nanorods were prepared using a seed-mediated growth method adapted from a previously reported protocol.^27^ All chemicals were obtained from Sigma-Aldrich and used without further purification.

The seed solution for gold nanorod growth was prepared as follows: 5 mL of 0.5 mM HAuCl₄ was combined with 5 mL of 0.2 M cetyltrimethylammonium bromide (CTAB) in a 20 mL vial. Fresh 0.01 M NaBH₄ (0.6 mL, diluted to 1 mL with water) was rapidly injected into the Au(III)-CTAB solution under vigorous stirring at 1,200 rpm. Stirring was stopped after 2 min, and the seed solution was aged at room temperature for 30 min.

For the growth solution, CTAB (43 mM) and NaOL (9 mM) were dissolved in 475 mL of water, followed by the addition of 12 mL of 4 mM AgNO₃ and 10 mL of 25 mM HAuCl₄. The pH was adjusted with 12 M HCl (1.5 mL or 2.1 mL for the two nanorod batches), and the solution was reduced using 0.8 mL of 100 mM ascorbic acid. The seed solution was then added to the growth solution at 30 °C and left undisturbed for 12 h to yield Au nanorods. Nanorods were collected by centrifugation and washed twice. As-synthesized nanorods were stabilized with 6 mM CTAB.

Washed nanorod resonance wavelengths were determined via ultraviolet-visible spectroscopy (Cary 6000i UV-Vis-NIR; Agilent Technologies). Size distributions were measured manually in FIJI, image analysis software, using transmission electron microscope images acquired at 200kV (Tecnai G2 F20 X-TWIN; Thermo Fisher Scientific).^50^ Gold concentrations were quantified by inductively coupled plasma optical emission spectroscopy (Thermo iCAP 6300; Thermo Fisher Scientific). Two nanorod batches were used in this study (Supplementary Fig. 1). Nanorods used for donors DN71, DN74, and DN76 exhibited a longitudinal resonance at 686 nm, with an aspect ratio of 2.73 ± 0.66 and a gold concentration of 420 µg/mL. Nanorods used for all other donors exhibited a resonance at 678 nm with an aspect ratio of 2.13 ± 0.78, and gold concentration of 350 µg/mL.

### Cell and nanorod sample preparation

Spectra were collected from T cells on day 13 or 14 post-CD3/CD28 activation. For JeKo-1 cells, measurements were collected 3-4 days after thawing. For primary B cells and patient PBMCs, measurements were collected on the day of thawing. Cells were washed three times in PBS and counted using an automated cell counter (LUNA-BX7™; Logos Biosystems). For SERS measurements, cells were mixed 1:1 by volume with gold nanorods (omitted for non-SERS controls) to generate final cell concentrations of approximately 6 × 10^4^ cells/µL (primary B and T cells and PBMCs), 2.1 × 10^4^ cells/µL (JeKo-1 B cells), and 1.5 × 10^5^ cells/µL (RBCs), sufficient to form an approximately single-cell layer at the base of the well. For co-culturing experiments, T cells and CD19-positive JeKo-1 B cells were mixed at a 3:1 effector-to-target ratio, common for *in vitro* assays that enrich for activation events.^41^ This is elevated from the physiological ratio observed *in vivo*, but allowed us to capture a high population of activated cells in our excitation spot.^51^

CTAB-stabilized nanorods were sonicated, then washed twice by centrifugation in water to remove excess cytotoxic surfactant^52^ while maintaining colloidal stability. After the final wash, nanorods were resuspended in half the original volume by sonication and mixed 1:1 with cells, restoring to the nominal stock concentration. Because washing introduces loss through aggregation and adhesion, the effective concentration is likely lower than nominal, but our protocol was kept consistent across all experiments.

### Liquid well preparation

Liquid wells were fabricated on 25 x 25 x 1 mm glass chips. Slides were plasma cleaned for 5 minutes at 100 W (PX-250 Plasma Asher; March Instruments; 85 mTorr, 2 SCCM O_2_, direct) and coated with a 5 nm Ti adhesion layer followed by 195 nm Au to provide reflectivity and low Raman background. Liquid wells were assembled following a method adapted from previous work.^53^ In brief, two layers of double-sided tape were hole-punched to create a spacer and applied to the gold-coated surface. The inner edges of the hole were coated in high-vacuum grease (Dow Corning) to prevent leakage. A 4 µL sample was loaded into the well and sealed with a borosilicate glass coverslip (Corning) to prevent sample evaporation during Raman acquisition.

### Flow cytometry / Fluorescence-Activated Cell Sorting

For activation analysis studies, CD19-CAR T cells, Mock T cells, and JeKo-1 B cells were washed by centrifugation and resuspended in phosphate buffered saline (PBS; Gibco). T cells were co-cultured with JeKo-1 B cells at a 3:1 effector-to-target ratio in PBS. For each donor, samples were collected at predefined timepoints (DN7518: 0, 25, 60, 90 min; DN2058 (in duplicate): 0, 25, 45, 65, 90 min). At each time point, an aliquot of ∼10^5^ cells was stained on ice with huCD3 (FITC; BioLegend; 300405), huCD19 (APC; BioLegend; 302211) at a 1:200 dilution, fixed (Fixation Buffer; Invitrogen), washed, and resuspended in flow buffer (PBS, 2% FBS, and 1 mM EDTA; Invitrogen). Single-color and unstained controls were used for compensation and gating (gating and all density plots are shown in Supplementary Fig. 12). Flow cytometry was performed on an ACEA Novocyte Quanteon (Agilent), and activation was tracked as CD3+CD19+ events using FlowJo software (BD Biosciences).^54^

For experiments using purified CD19-CAR T cells, DyLight 650-conjugated FMC63 anti-idiotype was used to label CD19-CAR T cells, which were then bulk sorted using a FACSAria II (BD Biosciences). Sorted cells were incubated overnight at 37°C and washed three times in PBS to allow fluorescent label removal before Raman analysis.

For mitochondrial mass measurements, donor-matched FMC-CAR, GD2-CAR, and Mock T cells were stained at room temperature for 20 minutes in the dark with a DyLight 650-conjugated anti-idiotype antibody to identify CAR+ and CAR-populations and with 2 µg/mL nonyl acridine orange (NAO; Invitrogen; A1372) to assess mitochondrial mass. Flow cytometry was performed on a Cytek Aurora and analysis was performed in FlowJo. Cells were gated for lymphocytes, singlets, and CAR expression (Supplementary Fig. 14a-c). NAO fluorescence was quantified as the median fluorescence intensity within the CAR+ and CAR-gates for CAR-transduced samples and across the full Mock population (Supplementary Fig. 14d-e).

For patient population FACS isolation, PBMCs were stained at room temperature for 20 minutes in the dark with anti-idiotype antibody (FITC; ACROBiosystems; FM3-FY45), 7-AAD (BioLegend; 420404), huCD3 (BV421; BioLegend; 317344), CD19 (PE; BioLegend; 302208), CD14 (PE; BioLegend; 399203), and CD235a (PE; BioLegend; 379009) at a 1:200 dilution, washed, and resuspended in flow buffer. Single-color and unstained controls were used for compensation and gating (gating and all density plots are shown in Supplementary Fig. 15a-b). Cells were sorted into CD3+CAR+, CD3+CAR-, and Dump+ populations. Dump+ cells were defined as PE+ cells labeled by a lineage dump channel containing CD19, CD14, CD235a antibodies. Sorting was performed on an FACSAria Fusion (BD Biosciences), and CAR+ T cell frequency was analyzed using FlowJo software.^54^

### CyTOF data analysis

Matched CyTOF data from the same patient cohort and timepoints were generated in a prior study and analyzed in this study as orthogonal validation of immune cell state.^48^ Analysis was done using FlowJo software^54^ and reported in Fig. 6b and Supplementary Table 5. The gating strategy is shown in Supplementary Fig. 15c. Cells were gated based on bead distance, event length, and DNA content. Single live cells were gated by excluding apoptotic cPARP+ events. Lymphocytes were selected based on CD45 expression. T cells were gated as CD3+ events and then analyzed for T cell state markers, including CAR expression, CD4, CD8α, Ki-67, CD57, T-bet, CD39, and PD-1.

### Mitochondrial extraction

Mitochondria were extracted from T cells using a mitochondria isolation kit for cultured cells (Thermo Scientific) according to the manufacturer’s protocol to obtain cytosolic and mitochondrial fractions. Mitochondrial pellets were lysed in 2% CHAPS (Abcam) in Tris-buffered saline. Mitochondrial-fraction protein concentration was quantified using a Bradford assay (Bio-Rad) by measuring absorbance at 595 nm on a spectrophotometer (Bio-Rad). For CAR-transduced samples, concentrations were adjusted by transduction efficiency to provide an estimated CAR+-adjusted mitochondrial protein concentration (assuming the non-transduced component resembles Mock).

### Raman spectroscopy

Raman spectra were collected on an XploRA+ confocal Raman microscope (HORIBA Scientific) equipped with a 1,200 gr/mm grating, 100 µm slit, 300 µm pinhole, and 785 nm diode excitation laser. Spectra were acquired over the 700-1,700 cm^−1^ biological fingerprint region for all donor-derived samples and 700-1,680 cm^−1^ for clinical samples on a rectangular scan grid with ∼30 µm spacing, exceeding both the laser spot size (∼1.2-1.5 µm) and typical cell diameters (7-12 µm), ensuring approximately single-cell sampling. All SERS measurements were taken with 1.5-second acquisition times.

Non-enhanced spectra (Supplementary Fig. 8) were obtained with 25-second acquisitions. Because of laser-induced cell drift (likely due to local heating) during longer exposures, spectra were collected in small batches (10-15 spectra) with the excitation location selected via point-by-point to remain centered on single cells.

Measurements for Figs. 3 and 5 were performed using a 50x/0.65 NA objective (Olympus LCPLAN 50X IR; Evident Scientific) with the correction collar set to 0.5. Under 785 nm excitation, this results in a calculated diffraction-limited spot size of ∼1.5 µm. Co-cultured activation and immune cell type measurements (Fig. 2 and 4) were taken using a 20x/0.80 NA objective (Olympus UPlanXApo; Evident Scientific), with a theoretical spot size of ∼1.2 µm. Actual spot sizes were larger due to sample-induced aberrations and scattering in liquid conditions. All spectra were acquired with 100% power output: 30.47 mW at the sample for the 20x objective and 26.34 mW for the 50x. Before acquisition, samples were allowed to settle for 15 minutes (co-culturing experiments) or 20 minutes (for single-cell experiments).

### Cell viability

CAR T cells were loaded into liquid wells described above, where tape is coated with high-vacuum grease to allow repeated opening for sample access. Cell viability was assessed using acridine orange/propidium iodide staining (Logos Biosystems) and quantified on an automated cell counter. Cells were incubated for 1 hour at room temperature under four conditions: with gold nanorods, without gold nanorods, with continuous 785 nm Raman excitation, and without laser exposure. Controls were also obtained at 0 minutes, with and without nanorod addition. Nanorod concentrations and Raman excitation were conducted in the same conditions used for SERS experiments. Each condition was performed in duplicate using independently prepared cell and nanorod samples.

### Scanning electron microscopy (SEM)

Cells were fixed in 4% paraformaldehyde (Sigma-Aldrich) and 0.1% glutaraldehyde (Sigma-Aldrich) in PBS for 30 min, followed by three washes to remove salts (one wash in PBS and two in water, with 5 min equilibration between spins). Fixed cells were mixed with nanorods and 2 µL droplets of the suspension were air dried on silicon chips. Samples were sputter-coated with ∼5 nm of Au/Pd alloy (60/40 wt%; Cressington 108 Auto Sputter Coater; Cressington Scientific Instruments). Imaging was performed on a FEI Magellan 400 XHR scanning electron microscope (Thermo Fisher Scientific) using the concentric backscattered detector at 5 kV landing voltage, 50 pA beam current, and −3 kV stage bias.

### Cryo-electron microscopy and tomography (cryo-EM/ET)

300 mesh molybdenum lacey carbon grids (LC300-MO; Electron Microscopy Sciences) were glow discharged for 10 seconds at 10 mA (PELCO easiGlow™ Glow Discharge Cleaning System; Ted Pella, Inc.). Cell-nanorod mixtures (3µL) were applied to each grid, blotted for 3 seconds, and vitrified in liquid ethane using a Vitrobot^TM^ Mark IV (Thermo Fisher Scientific). Grids were imaged on a Glacios^TM^ 200kV cryo-transmission electron microscope using Tomography 5 (Thermo Fisher Scientific).

Tomographic tilt series were acquired from −60° to +60°. Tilt images were aligned and reconstructed using IMOD (Etomo interface).^55^ Reconstructed tomograms were visualized in UCSF ChimeraX^56^.

### ζ-potential characterization

Cell and nanorod ζ-potential values were measured in a 1:1 mixture of PBS and water to match experimental conditions using Phase Analysis Light Scattering (Nanobrook Omni; Brookhaven Instruments). ζ-potential measurements were performed for mammalian immune cells, red blood cells, and bacterial reference strains to benchmark relative surface charges. Each sample was measured in triplicate, with 30 cycles per measurement. Red blood cells, which have more heavily sialylated membrane glycoproteins, exhibited a more negative ζ-potential (∼ −17 mV) than white blood cells (∼ −12 mV).^57^ This was consistent with the greater nanorod binding observed in dried SEM samples (Fig. 2e-f).

### Single-cell RNA sequencing analysis

Single-cell RNA sequencing (scRNA-seq) was performed on CD19 CAR T cell infusion products from seven patients with large B-cell lymphoma (LBCL) treated with axicabtagene ciloleucel (axi-cel; Yescarta, Kite Pharma) as standard-of-care therapy at our institution. Patients provided informed consent through Clinical Outcomes Biorepository (Stanford IRB #43375), and clinical metadata were obtained via retrospective chart review.

Raw sequencing data were processed and analyzed using Seurat v5.0.^58^ Cells were filtered to remove ones with greater than 10% mitochondrial gene content, as well as low-quality cells based on total UMI counts and number of detected genes. After quality control, the final dataset comprised 102,359 cells across 36,604 genes.

To identify transcriptional differences between CD19 CAR-expressing (CAR+; n = 82,552) and non-expressing (CAR-; n = 19,807) cells, differential expression analysis was performed using Seurat’s FindMarkers pipeline. Only genes expressed in ≥5% of cells in either group were retained. TCR and BCR clonotypes, as well as sex-associated genes, were removed prior to testing. The Wilcoxon rank-sum test was used as the default statistical method to identify differentially expressed genes (DEGs) and differential regulon activity and results were visualized using the EnhancedVolcano package.

Transcription factor network (regulon) activity was inferred using SCENIC (Single-Cell rEgulatory Network Inference and Clustering).^59^ Regulons were scored per cell using the AUCell algorithm,^60^ and differential regulon activity between CAR+ and CAR-cells was assessed using the Wilcoxon rank-sum test. Pathway enrichment analysis was performed on differential expression analysis. Adjusted p-values were computed using the Benjamini–Hochberg procedure to control the false discovery rate.

### Data preprocessing

All preprocessing and computational analyses were performed in Jupyter notebooks (Python 3.11.13) and executed on a CPU.

Raman spectra were preprocessed following a method adapted from previous work that included despiking, denoising, baseline correction, and normalization (Supplementary Fig. 4).^23^ Cosmic ray spikes were removed using a modified Whitaker-Hayes z-score method.^61^ Despiked spectra were denoised by wavelet thresholding (BayesShrink; skimage) to suppress high-frequency noise while preserving Raman peaks.^62^ Baseline drift was corrected using adaptive iteratively reweighted penalized least squares (airPLS).^63^ Finally, spectra were z-score standardized on a per-spectrum basis for cross-sample comparisons and downstream analyses. For each acquisition period, spectra with average unnormalized intensities exceeding two standard deviations above that experiment’s mean were excluded to avoid laser-induced artifacts or detector saturation. Before analysis, spectra were linearly interpolated onto a common wavenumber axis across the acquired spectral range (700-1,680 cm^−1^ for patient spectra and 700-1,700 cm^−1^ for donor-derived spectra) to ensure equal feature dimensionality across compared datasets.

A subset of patient Raman maps (indicated with an asterisk in Supplementary Table 6) had acquisition fields that partially overlapped the liquid well edge. These maps were noted during acquisition and confirmed using spatial intensity maps (mean unnormalized intensity plotted as a function of coordinate). For these maps, off-well regions were excluded from downstream analysis using coordinate-based masks, initially guided by contiguous low-intensity regions at the map boundary and manually adjusted when necessary to fully capture the visible off-well area. Final exclusions were applied using acquisition coordinates and were defined without reference to sample type or downstream classification results.

### Spectral plotting and intensity analyses

To uncover spectral differences between conditions, we computed per-donor intensity difference curves by subtracting class-averaged spectra. Difference curves were normalized to range −1 to 1, then median-aggregated across donors and plotted with 95% confidence bands. To analyze time-resolved activation dynamics, spectra were grouped into time bins labeled by their earliest time point (15-25, 25-35, 35-45, 45-55, 55-75, and 75-95 min). For plotting, mean spectra were computed within each time bin. Temporal changes were tracked in 20 wavenumber bands centered on protein and aromatic signatures (898, 1,002, 1,025, 1,222, 1,262, 1,602 cm^−1^)^30,33,35^ and nucleic acid signatures (728, 1,092, 1,334 cm^−1^).^20,29–31,33,36^ For activation analysis, spectral intensities in target-engaged CAR T cell co-cultures were computed relative to Unstim controls using log-ratio differences to remove global acquisition-related variation with time. Within each biochemical class, intensities were z-score standardized within bands and averaged across them. Confidence intervals were estimated using block-bootstrap resampling. Monotonicity of trends was assessed using two-sided Spearman’s rank correlation between z-scored intensities and experimental time (scipy).

For patient cohort biochemical band analysis, timepoint changes were tracked using 20 cm^−1^ wide windows centered at nucleic acid-associated signatures (728, 1,092, 1,334 cm^−1^)^20,29–31,33,36^ and protein and aromatic-associated bands (898, 1,002, 1,025, 1,251, 1,583, and 1,645 cm^−1^).^29,30,33,35^ For each spectrum, the NA-protein spectral contrast was computed as the mean intensity across nucleic acid-associated bands minus the mean intensity across protein-associated windows. Timepoint-specific changes were then computed by subtracting the matched Pre mean contrast from each timepoint contrast, forming the NA-protein spectral contrast plotted in Fig. 6e. Confidence intervals were estimated by bootstrap resampling of single-cell spectra within each patient/timepoint group.

### Dimensionality reduction and manifold projection

Preprocessed spectra were visualized using Uniform Manifold Approximation and Projection (UMAP; n_components = 2, n_neighbors = 20, min_dist = 0.1, metric = “manhattan”).^28^ UMAPs were generated using spectra balanced by donor and class to avoid overweighting. For qualitative illustration of activation dynamics, we computed the centroid of each class within each time bin by calculating the mean UMAP coordinate. Uncertainty in centroid location was estimated using the standard error of the coordinates.

### Machine learning classification and feature importance

Lightweight ensemble Light Gradient Boosting Machine (LightGBM) models were trained for spectral classification.^64^ LightGBM demonstrated the best trade-off between accuracy and computational efficiency in our application among the ensemble models evaluated (Supplementary Table 3). Hyperparameter tuning and overfitting control were performed using only the training data. First- and second-order Savitzky-Golay derivatives (window_length = 11, polyorder = 3; scipy) were appended as additional dimensions so that slope and curvature information could also inform classification accuracy.^65^ Derivative-augmented data were used for all classification analysis; non-augmented spectra were used for feature-importance analysis to preserve interpretability. For clinical sample analysis, classification models were trained only on datasets with sufficient spectral counts, defined as at least 250 spectra per class or condition (Supplementary Table 6).

Prior to model training, datasets were split into 80:20 train:test sets, stratified by class. Hyperparameters were optimized via Bayesian optimization (Optuna)^66^ using five-fold cross-validation (CV) stratified by class, with early stopping based on validation performance (training terminated if AUC didn’t improve for 150 consecutive boosting rounds), and a maximum of 2,000 boosting rounds. The scale_pos_weight parameter was adjusted based on class imbalance. Learning curves from five-fold CV were used to estimate the optimal number of boosted trees (n_estimators) that avoided overfitting while maximizing validation AUC. Final models were retrained and evaluated using 10-fold CV stratified by class on the training data, and performance was assessed through confusion matrices (scikit-learn). Receiver operating characteristic (ROC) curves and their AUC values were computed on held-out test sets (scikit-learn). For per-donor models, ROC curves were interpolated onto a common grid, bootstrap averaged, and plotted with 95% confidence intervals.

For immune cell type classification and mixed patient PBMC timepoint classification, both of which were multiclass tasks, the same general workflow was used, except that hyperparameters were optimized using log loss minimization rather than AUC maximization. For CAR versus Mock T cell classification, when models were trained across all donors, pooled accuracy reached 69.3% for CD19-CAR versus Mock and 81.4% for GD2-CAR versus Mock. However, UMAP visualization showed that inter-donor variability exceeded between-class differences (Supplementary Fig. 6a), indicating a strong donor prior and motivating a donor-level training scheme, given the available dataset sizes.^67^ Patient multiclass and binary classifications were similarly performed using patient-level or patient/timepoint grouping-level LightGBM models. Multiclass PBMC timepoint classification performance was reported as balanced accuracy for each CV fold. CD3+CAR+ versus CD3+CAR-T cell and CD3+CAR+ versus non-CAR background binary classification performance was reported as both accuracy and AUC per fold.

To identify spectral features that best improved our model’s accuracy, we used LightGBM’s intrinsic gain-based feature importance scores for models trained on non-derivative-augmented spectra.^64^ Per-donor importance profiles were sum-normalized to account for magnitude variation and scaled from 0 to 1. Mean gain profiles were then computed, and uncertainty was estimated using the standard deviation across donors.

### CAR-associated Raman score analysis

Patient-specific binary LightGBM classifiers were trained using pre-infusion mixed PBMC spectra as the CAR-negative reference class and flow-isolated CD3+CAR+ spectra collected post-infusion as the CAR-positive reference class. Post-infusion CD3+CAR+ reference spectra were included only from patient/timepoint groups meeting the spectral-count inclusion threshold (see Data size and handling). For Patient 022, Peak and Late CD3+CAR+spectra were pooled for training. For Patients 019 and 024, only Late CD3+CAR+ spectra were used because of insufficient Peak CD3+CAR+ spectral counts. Patient 016 was excluded because of overall insufficient CD3+CAR+ spectral counts. After training, each classifier was then applied to matched Peak and Late mixed PBMC spectra from the same patient. The classifier output probability for the CAR-positive class was defined as the CAR-associated Raman score. Mean Raman scores were computed for each patient and compared with matched flow-measured CAR+ T cell frequencies.

### Pseudotime analysis

Pseudotime trajectories were generated from spectra projected into a 45-dimensional PCA space (scikit-learn). We computed a minimum spanning tree, rooted at the earliest time point cell, on the largest connected component of a k-nearest-neighbor graph (k-NN, k = 30 for DN2058, k = 40 for DN7518; cosine distance). For trajectory visualization (Supplementary Fig. 11), pseudotime values were binned into 12 clusters, and the centroid of each cluster was overlaid onto a UMAP of PCA-reduced data. Correlation between pseudotime and experimental time was evaluated using two-sided Spearman’s rank correlation (scipy).

### Data size and handling

The number of independent single-cell Raman spectra collected for each donor, patient, and condition is reported in Supplementary Tables 2 and 6. Across donor-derived experiments, sample sizes ranged from 804 to 6,189 spectra per condition and donor. For patient-derived experiments, spectral counts ranged from 37 to 1,069 spectra per condition and patient, with downstream classification restricted to datasets containing at least 250 spectra. Clinical datasets with fewer than 250 spectra are reported in Supplementary Table 6 but considered insufficient for further analysis. Each spectrum corresponds to a distinct single cell; no repeated measurements of the same cell were taken. All analyses were performed per donor, per patient, or per patient/timepoint grouping, as appropriate, using stratified cross-validation when applied for classification models. No statistical models including covariates were used.

## Data Availability

The source data that support the main findings in this work will be deposited in a public repository and made available upon publication of the manuscript. Raw single cell RNA sequencing data are available through Gene Expression Omnibus as part of C Y Tsui et al. [in submission]. Raw CyTOF data from the matched patient cohort are available through the Stanford Digital Repository at https://purl.stanford.edu/qb215vz6111, originally published by Good et al.^48^

## Code Availability

The code used to analyze and classify Raman spectral data will be deposited in a public repository and made available upon publication of the manuscript. The workflow relies on established Python packages and does not introduce custom ML algorithms.

## Acknowledgements

A.S. acknowledges financial support from the National Science Foundation (NSF) through the Graduate Research Fellowships Program. B.O. acknowledges support from the Stanford Bio-X Bowes Graduate Student Fellowship. Y.L. was supported by the Stanford Science Fellowship. This work was in part supported by NIH Grants: R21CA301006 (J.A.D.) 2P01CA049605 (D.B.M., C.L.M., Z.G.), 1OT2OD038101 (C.L.M., Z.G.), R35CA283888 (C.L.M.), 2P30CA124435-16 (C.L.M.), the Virginia and D.K. Ludwig Fund for Cancer Research (C.L.M.), an EPICC Translational Research Grant (St. Baldrick’s Foundation, C.L.M.), the Kona Innovation Challenge Award (C-04134) from the Parker Institute for Cancer Immunotherapy (PICI) (Z.G.), PICI C-04248 (D.B.M.), Stanford Center for Digital Health Award (Z.G.), Laude Institute MOONSHOTS \\ ONE Honorable Mention Award (Z.G.), and a sponsored research agreement with Kite Pharma, a subsidiary of Gilead Sciences (D.B.M.). Z.G. was supported by the Parker Bridge fellowship (PICI C-02895) and the NIH/NCI Pathway to Independence award (1K99CA293149, 4R00CA293149). Z.G. and C.L.M. are members of the Parker Institute for Cancer Immunotherapy, which supports the Stanford University Cancer Immunotherapy Program. Z.G., C.L.M., D.B.M., and E.S. are Weill Cancer Hub West Investigators (Team PROMISE). J.A.D is a Chan Zuckerberg Biohub – San Francisco Investigator and acknowledges funding from the Chan Zuckerberg Biohub San Francisco. Select elements of Figures 1, 3, 4, and 5 were adapted from Biorender.com. TEM/SEM imaging, zeta potential, gold coating, and Raman spectroscopy was performed at nano@stanford RRID:SCR_026695. From these facilities we would like to thank Dr. Christina Newcomb for training and discussions on Raman acquisition, Tom Carver for gold-coated slide preparation, and Richard Chin for providing training in SEM imaging. Cell sorting and flow cytometry analysis for this project was done on instruments in the Stanford Shared FACS Facility (RRID: SCR_017788). Cryo-EM/ET were performed at the Stanford University Cryo-electron Microscopy Center (cEMc). The authors acknowledge the use of facilities and technical assistance of the Stanford Doerr School of Sustainability Environmental Measurements Facility (RRID: SCR_023255). Nanoparticle concentrations were measured by Dr. Guangchao Li at the Stanford Environmental Measurements Facility. Additionally, we would like to thank Liam Herndon for preparing samples for gold concentration quantification, Dr. Jason Casar for manuscript feedback, and Drs. Maia Stiber, Michael Stiber, and Nhat Vu, for insights on machine learning analysis. We are grateful to members of the Appel laboratory for their discussions and support and to members of the Dionne, Good, and Mackall laboratories for their collaboration throughout this work. This paper has been proofread by a language model (AI). The authors have consulted the output, edited the paper wherever deemed appropriate, and are responsible for the content. The content is solely the responsibility of the authors and does not necessarily represent the official views of the NIH.

## Author Contributions

All authors contributed to the review and editing of the manuscript text and figures. A.S., C.L.M., Z.G., and J.A.D. conceived of and designed the experiments. A.S. led experimental and computational work, including cell culture, sample preparation, nanorod and cell characterization (electron microscopy, UV-Vis, zeta potential), Raman spectral acquisition, flow cytometry, viability assays, CyTOF data analysis, data preprocessing, machine-learning analysis, and code implementation. A.S additionally drafted the figures and text of the manuscript. B.Q. contributed to biological experimental design, isolated and engineered T cell samples, and assisted with flow cytometry sorting, measurements, and analysis. B.O. assisted with early project conceptualization, development of experimental protocols, sample preparation, and spectroscopy measurements.

A.G. assisted with sample preparation, spectroscopy measurements, data analysis, and development of experimental protocols. K.C. contributed preprocessing code and provided input on computational structure and Raman analysis. Y.L. synthesized gold nanorods. P.Q. assisted with T cell sample isolation and engineering. H.W. assisted with cryo-EM/ET imaging and tomogram reconstruction. K.C.Y.T. assisted in the analysis and creation of the data, figure, and text associated with Supplementary Fig. 7. C.A. assisted with sample preparation and Raman acquisition. E.S. contributed to experimental design. D.B.M. provided clinical samples used in this study. A.S., B.Q., E.S., Z.G., and J.A.D. contributed to the biological interpretation of the results. C.L.M., Z.G., and J.A.D. conceived the idea, supervised the work, and provided materials, reagents, and expertise necessary in immunology, plasmonics, microscopy, spectroscopy, and data analysis. All authors approved the final manuscript.

## Additional Information

Supplementary Information is available for this paper.

**Supplementary Video 1:** Aligned cryo-ET tilt series of a whole T cell with gold nanorods Aligned cryo-electron tomography (cryo-ET) tilt series of a T cell with gold nanorods deposited on a lacey carbon grid (corresponding in Supplementary Fig. 2a). Smaller high-contrast features represent nanorods present on the cell surface and grid.

**Supplementary Video 2:** Reconstructed tomogram of the tilt series shown in Supplementary Video 1 Three-dimensional tomogram reconstructed from the aligned tilt series shown in Supplementary Video 1. High-intensity features correspond to nanorods.

**Supplementary Video 3:** Aligned cryo-ET tilt series of two T cells with gold nanorods Aligned cryo-electron tomography (cryo-ET) tilt series of two T cells with gold nanorods deposited on a lacey carbon grid (corresponding in Supplementary Fig. 2b). Smaller high-contrast features represent nanorods present on the cell surface and grid.

