## Supplementary Info for "Dynamic, single-cell monitoring of CAR T cell identity and activation with Raman spectroscopy"

### Table of Contents

SUPPLEMENTARY TABLE 1: TRANSDUCTION EFFICIENCIES FOR ALL DONORS 3
SUPPLEMENTARY FIG. 1: GOLD NANOROD CHARACTERIZATION 4
SUPPLEMENTARY FIG. 2: CRYO-ELECTRON MICROGRAPHS OF T CELLS WITH NANORODS 5
SUPPLEMENTARY FIG. 3: CELLULAR SERS VS. NANOROD AND NON-SERS CONTROLS 6
SUPPLEMENTARY TABLE 2: SPECTRAL COUNT FOR HEALTHY DONOR AND PRECLINICAL EXPERIMENTS 7
SUPPLEMENTARY FIG. 4: RAMAN SPECTRAL PREPROCESSING WORKFLOW 10
SUPPLEMENTARY TABLE 3: BENCHMARKING OF ENSEMBLE CLASSIFIERS FOR RAMAN SPECTRA 11
SUPPLEMENTARY FIG. 5: IMMUNE CELL TYPE SPECTRAL ANALYSIS 12
SUPPLEMENTARY TABLE 4: SERS BAND ASSIGNMENTS FOR IMMUNE CELL SPECTRA 13
SUPPLEMENTARY FIG. 6: PRE-SORTED CD19-CAR SPECTRAL ANALYSIS ACROSS DONORS 15
SUPPLEMENTARY FIG. 7: TRANSCRIPTOMIC SIGNATURES OF CAR+ VERSUS CAR- CELLS IN CD19 CAR T INFUSION PRODUCTS 16
SUPPLEMENTARY FIG. 8: NON-SERS CD19 CAR VS MOCK RAMAN DATA 17
SUPPLEMENTARY FIG. 9: LOSS OF ANTI-IDIOTYPE LABEL AFTER POST-SORT INCUBATION 18
SUPPLEMENTARY FIG. 10: POPULATION-LEVEL CO-CULTURE RAMAN SIGNATURES 19
SUPPLEMENTARY FIG. 11: PSEUDOTIME TRAJECTORY ANALYSIS AND CORRELATION WITH REAL TIME 20
SUPPLEMENTARY FIG. 12: FLOW CYTOMETRY CHARACTERIZATION OF ANTIGEN-SPECIFIC ACTIVATION 21
SUPPLEMENTARY FIG. 13: GD2-CAR AND MOCK SERS SPECTRAL CLASSIFICATION ACROSS DONORS 23
SUPPLEMENTARY FIG. 14: REPRESENTATIVE MITOCHONDRIAL MEASUREMENTS IN FMC-CAR, GD2-CAR, AND MOCK T CELLS 24
SUPPLEMENTARY TABLE 5: EXPANDED FLOW CYTOMETRY AND CYTOF VALIDATION VALUES FOR LONGITUDINAL PATIENT SAMPLES 25
SUPPLEMENTARY FIG. 15: REPRESENTATIVE GATING FOR FLUORESCENCE-ACTIVATED CELL SORTING (FACS) AND MASS CYTOMETRY (CYTOF) ANALYSIS OF LONGITUDINAL PATIENT SAMPLES 28
SUPPLEMENTARY TABLE 6: SPECTRAL COUNT FOR LONGITUDINAL PATIENT SAMPLES (FIGURE 6) 30
SUPPLEMENTARY FIG. 16: LONGITUDINAL MIXED PBMC CLINICAL SAMPLE ANALYSIS ACROSS PATIENTS 32
SUPPLEMENTARY FIG. 17: LONGITUDINAL PBMC BIOCHEMICAL BAND INTENSITY TRENDS 33
SUPPLEMENTARY FIG. 18: LONGITUDINAL FLOW-ISOLATED CLINICAL SAMPLE ANALYSIS ACROSS PATIENTS INCLUDED IN FIG. 6E 35
SUPPLEMENTARY FIG. 19: CLINICAL SAMPLE CAR-ASSOCIATED RAMAN SCORE FOR PATIENTS INCLUDED IN FIG. 6F 36
REFERENCES 38

**Supplementary Table 1:** Transduction efficiencies for all donors

| Donor ID | CAR construct | Transduction efficiency (%) |
|---|---|---|
| DN71 | CD19-CAR | N/A |
| DN74 | CD19-CAR | 96.1 |
| DN76 | CD19-CAR | 96.0 |
| DN81 | CD19-CAR | 76.0 |
| DN2058 | CD19-CAR | 71.7 |
| DN4400 | CD19-CAR | 78.3 |
| | GD2-CAR | 58.6 |
| DN4402 | CD19-CAR | 54.9 |
| | GD2-CAR | 41.2 |
| DN4408 | CD19-CAR | 55.8 |
| | GD2-CAR | 34.8 |
| DN4411 | CD19-CAR | 58.1 |
| | GD2-CAR | 41.4 |
| DN4414 | CD19-CAR | 59.1 |
| | GD2-CAR | 39.0 |
| DN7518 | CD19-CAR | 74.8 |

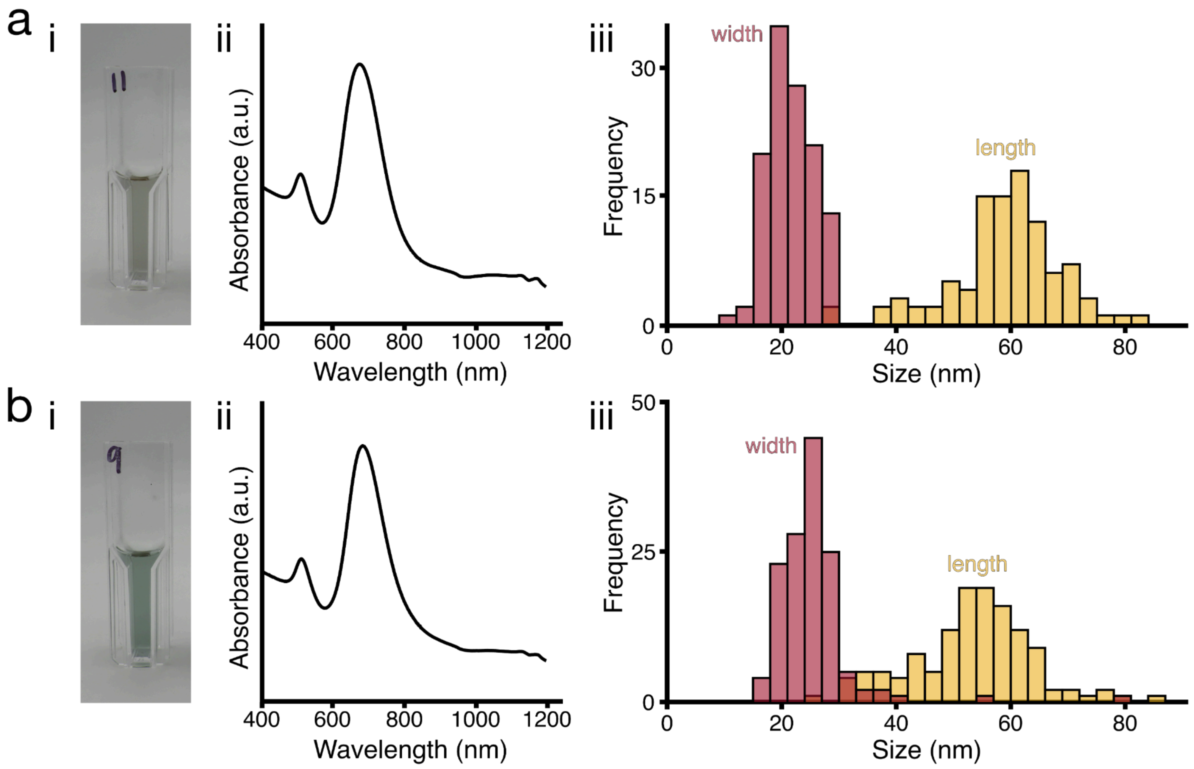


**Supplementary Fig. 1:** Gold nanorod characterization

Images of diluted nanorod suspensions (i), extinction spectra (ii), and size distributions (iii) for gold nanorods used for **(a)** donors DN71, 74, 76 (Fig. 3) and **(b)** all other donors. Both samples display a weak transverse plasmon resonance at ~520 nm and a strong longitudinal resonance at ~680 nm. Nanorods in (a) have an aspect ratio of 2.73 ± 0.66, and those in (b) have an aspect ratio of 2.13 ± 0.78.

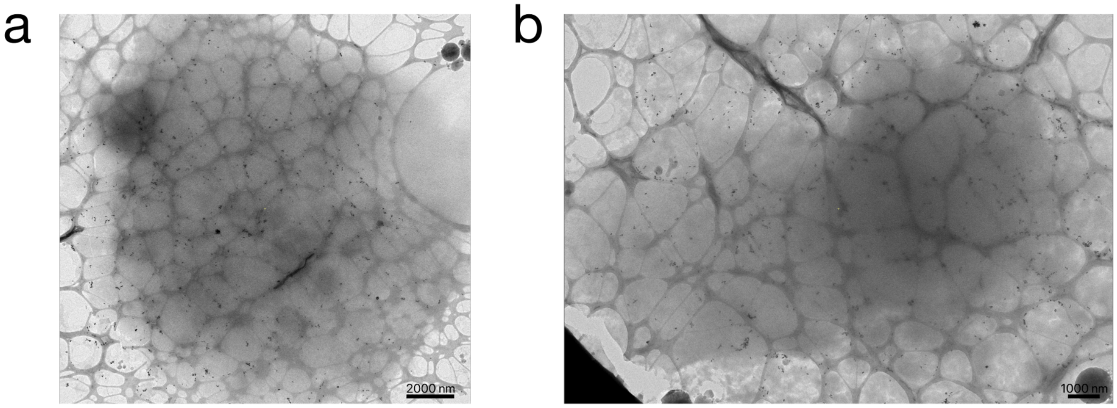


**Supplementary Fig. 2:** Cryo-electron micrographs of T cells with nanorods

Cryo-transmission electron micrographs of **(a)** a single frozen T cell and **(b)** two frozen T cells mixed with gold nanorods on lacey carbon grids. Smaller high-contrast features correspond to nanorods present on the cell surface and the grid. Supplementary Videos 1 and 3 show aligned cryo-tomography tilt series of the cells in (a) and (b), respectively, and Supplementary Video 2 includes a reconstructed tomogram of the cell in (a).

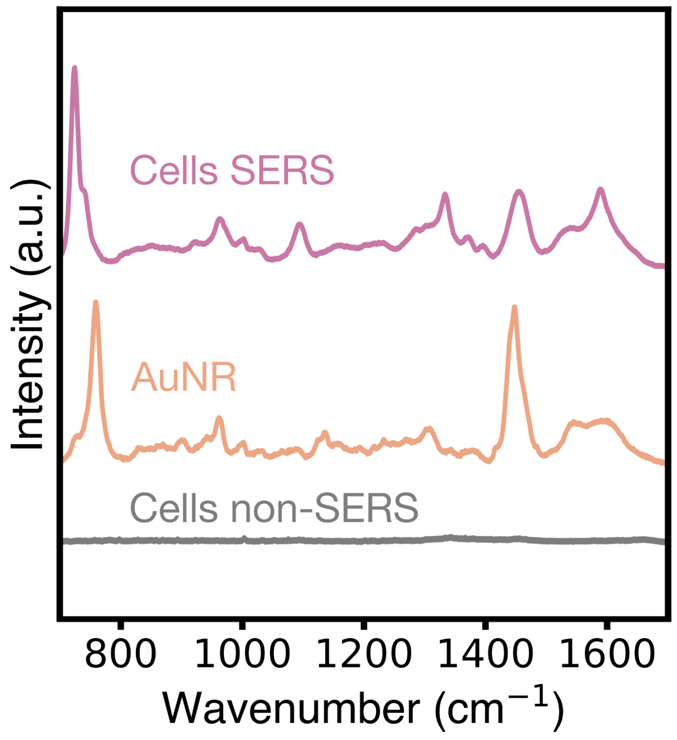


**Supplementary Fig. 3:** Cellular SERS vs. nanorod and non-SERS controls

Unnormalized Raman spectra of gold nanorods (AuNR, orange), cells without nanorods (cells non-SERS, grey), and cells with nanorods (cells SERS, pink), all taken with the same acquisition parameters. "Cells SERS" spectra exhibit significantly higher signal intensity than "cells non-SERS" spectra. Peaks in the AuNR spectrum likely arise from cetyltrimethylammonium bromide (CTAB) surfactant. The sharp bands in the AuNR spectrum, particularly the prominent 770 and 1445 cm$^{-1}$ features, are substantially reduced in the cellular SERS spectrum, consistent with minimal ligand contribution relative to cellular biomolecular signals.

**Supplementary Table 2:** Spectral count for healthy donor and preclinical experiments

| Class | Donor ID | Spectra count |
|---|---|---|
| **Figure 2: Blood Cell Type** | | |
| T cells | DN7518 | 855 |
| | DN2058 | 1180 |
| Primary B cells | — | 1972 |
| JeKo-1 B cells | — | 4653 |
| Red blood cells | — | 2342 |
| **Figure 3: CD19-CAR vs Mock (unsorted)** | | |
| CD19-CAR | DN71 | 1988 |
| | DN74 | 3444 |
| | DN76 | 6151 |
| | DN81 | 1241 |
| | DN4400 | 1162 |
| | DN4402 | 1179 |
| | DN4408 | 1173 |
| | DN4411 | 1161 |
| | DN4414 | 1169 |
| Mock | DN71 | 1975 |
| | DN74 | 3158 |
| | DN76 | 6189 |
| | DN81 | 1254 |
| | DN4400 | 1177 |
| | DN4402 | 1182 |

| | | |
|---|---|---|
| | DN4408 | 1176 |
| | DN4411 | 1239 |
| | DN4414 | 1169 |
| **Figure 3: CD19-CAR vs Mock (sorted)** | | |
| CD19-CAR | DN4400 | 1176 |
| | DN4408 | 1172 |
| | DN4411 | 1167 |
| | DN4414 | 1552 |
| Mock | DN4400 | 1261 |
| | DN4408 | 1174 |
| | DN4411 | 1173 |
| | DN4414 | 1541 |
| **Figure 4: Co-cultured CD19-CAR vs Mock** | | |
| CD19-CAR T & JeKo-1 B cells | DN7518 | 2216 |
| | DN2058 | 1489 |
| Mock T & JeKo-1 B cells | DN7518 | 2239 |
| | DN2058 | 1509 |
| **Figure 4: GD2-CAR vs Mock** | | |
| GD2-CAR | DN4400 | 1177 |
| | DN4402 | 1184 |
| | DN4408 | 1171 |
| | DN4411 | 1178 |
| | DN4414 | 1155 |
| | DN4400 | 1177 |

| | | |
|---|---|---|
| Mock | DN4402 | 1182 |
| | DN4408 | 1176 |
| | DN4411 | 1239 |
| | DN4414 | 1169 |
| **Supplementary Figure 7: non-SERS CD19-CAR vs Mock** | | |
| GD2-CAR | DN71 | 804 |
| | DN74 | 1121 |
| | DN76 | 1838 |
| Mock | DN71 | 1044 |
| | DN74 | 1246 |
| | DN76 | 1755 |

**Note**: Spectral counts are of analyzed spectra post-filtering for laser-induced damage or detector saturation (Supplementary Fig. 4).

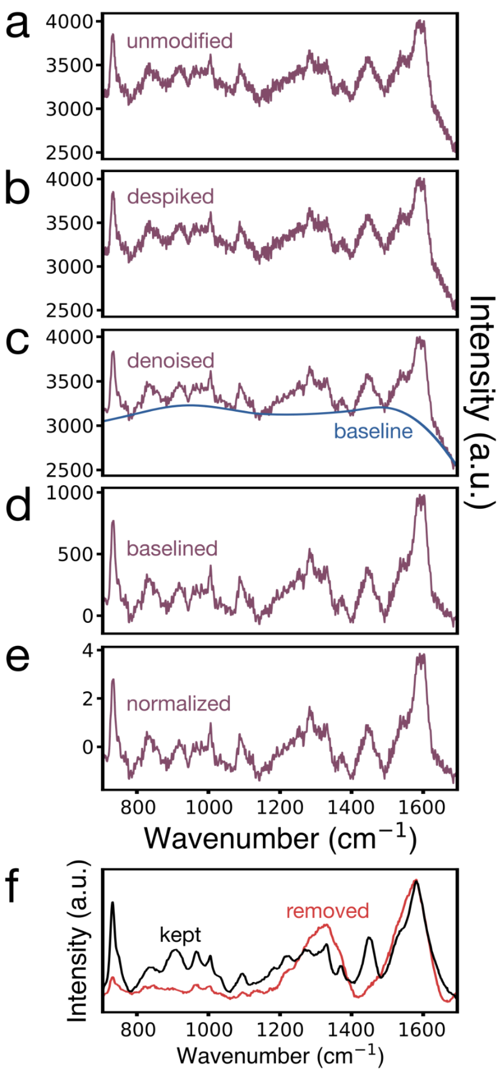


**Supplementary Fig. 4:** Raman spectral preprocessing workflow

A representative SERS spectrum of an immune cell shown **(a)** as acquired, **(b)** after cosmic ray removal,[1] **(c)** following wavelet-based denoising (baseline shown in blue),[2] **(d)** after adaptive iteratively reweighted penalized least squares baseline correction,[3] and **(e)** after normalization. **(f)** To exclude spectra impacted by laser-induced damage or detector saturation, spectra with average unnormalized intensities greater than 2 standard deviations above the mean for that experiment are removed. Shown are representative normalized average spectra retained (black) and removed (red) from one experiment.

**Supplementary Table 3:** Benchmarking of ensemble classifiers for Raman spectra

| Model | F1 | Accuracy | Train time (s) | Inference time (µs/sample) |
|---|---|---|---|---|
| Random Forest | 0.71 ± 0.02 | 0.72 ± 0.02 | 8.1 ± 3.4 | 27.2 ± 9.8 |
| Support Vector Machine | 0.67 ± 0.00 | 0.50 ± 0.00 | 17.2 ± 2.8 | 2148.5 ± 106.0 |
| Explainable Boosting Machine | 0.75 ± 0.00 | 0.76 ± 0.00 | 291.5 ± 4.6 | 97.1 ± 3.1 |
| CatBoost | 0.74 ± 0.00 | 0.75 ± 0.00 | 53.0 ± 29.2 | 868.4 ± 326.0 |
| XGBoost | 0.73 ± 0.00 | 0.74 ± 0.00 | 27.5 ± 0.9 | 15.2 ± 2.8 |
| LightGBM | 0.74 ± 0.00 | 0.74 ± 0.00 | 9.5 ± 0.5 | 12.2 ± 1.0 |

**Note**: For a controlled comparison across models, we evaluated performance on a single-donor CD19-CAR vs Mock preprocessed Raman dataset (~1000 spectra/class) with added first- and second-derivative dimensions. All models were trained on the same stratified 80/20 train/test split, hyperparameters were tuned using an equal-budget randomized search (20 trials/model), and performance (accuracy and F1-score) was evaluated on the held-out test set.

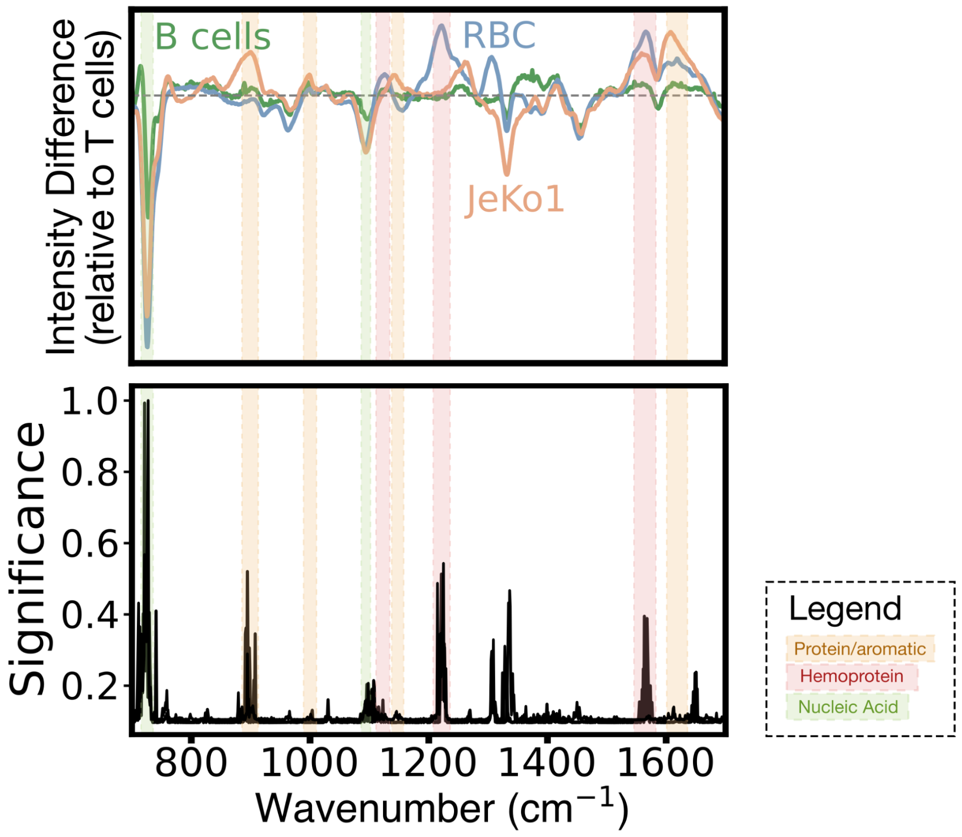


**Supplementary Fig. 5:** Immune cell type spectral analysis

Mean spectral difference of primary human B cells, JeKo-1 B cells, and red blood cells (RBC) relative to T cells (top). The grey dashed line marks zero difference. Feature importance plot showing the contribution of each spectral dimension (wavenumber) to classification among T cells, primary B cells, JeKo-1 B cells, and RBCs using a trained ML model (bottom). Colored spectral bands highlight protein/aromatic (gold), hemoprotein (red), and nucleic acid (green) regions.

**Supplementary Table 4:** SERS band assignments for immune cell spectra

| **Wavenumber ($cm^{-1}$)** | **Assignment** |
|---|---|
| 717-721 | Phosphocholine; symmetric C-N stretch[4] |
| 724 | Hypoxanthine; porphyrin ring mode[5] |
| 725-735 | Adenine; ring breathing mode[6–8] |
| 749-755 | Cytochrome c; heme breathing mode[9] |
| 870-880 | Phosphocholine; symmetric C-N stretch[4] |
| 880-900 | Tryptophan; indole ring mode[10,11] |
| 960-964 | Guanine[12] |
| 1,002-1,004 | Phenylalanine; ring breathing (C-C skeletal)[10,11] |
| 1,020-1,031 | Phenylalanine; C-N stretch[7] |
| 1,090-1,100 | Nucleic acid backbone; $PO_2^-$ symmetric stretch[6–9] |
| 1,127-1,129 | Hemoprotein; porphyrin ring mode[5] |
| 1,130-1,131 | Cytochrome c[9] |
| 1,140-1,160 | Protein; C-C and C-N stretch[6] |
| 1,175-1,177 | Aromatic amino acids (Tyr/Phe/Trp); C-H bending / scissoring[7,10] |
| 1,220-1,300 | Protein backbone; amide III region[10,11] |
| 1,225 | Deoxygenated hemoglobin[13] |
| 1,255-1,266 | α-helical protein; amide III region[10] |
| 1,310-1,311 | Cytochrome c[9] |
| 1,323-1,330 | Lipids; C-C, C-H stretches[9,11] |
| 1,330-1,346 | Nucleic acids (G/A/pyrimidines); ring C-N and C=N stretches[9,10,12] |
| 1,450-1,458 | Nucleic acids[12] |
| 1,455-1,465 | Lipids; $CH_2$/$CH_3$ deformation[6] |

| | |
|---|---|
| 1,480-1574 | Protein; amide II region[10,11] |
| 1,552-1,560 | Tryptophan; indole ring stretch[6,11] |
| 1,565-1,570 | Hemoprotein; porphyrin ring mode[5] |
| 1,584-1,590 | Cytochrome c[9] |
| 1,587-1,615 | Aromatic amino acids (Tyr/Phe/Trp); C=C/C=O stretches[7,10,11] |
| 1,620-1,625 | Hemoprotein; porphyrin ring $C_a$=$C_b$ C=C stretch[5] |
| 1,620-1,700 | Protein backbone; amide I region[10] |

**Note**: Raman peaks will vary based on cell type and culture conditions, SERS substrate (or lack thereof), and acquisition parameters such as excitation wavelength. Peak assignments should therefore be interpreted as approximate.

**Abbreviations:** Tyr: tyrosine; Phe: phenylalanine; Trp: tryptophan; G: guanine; A: adenine

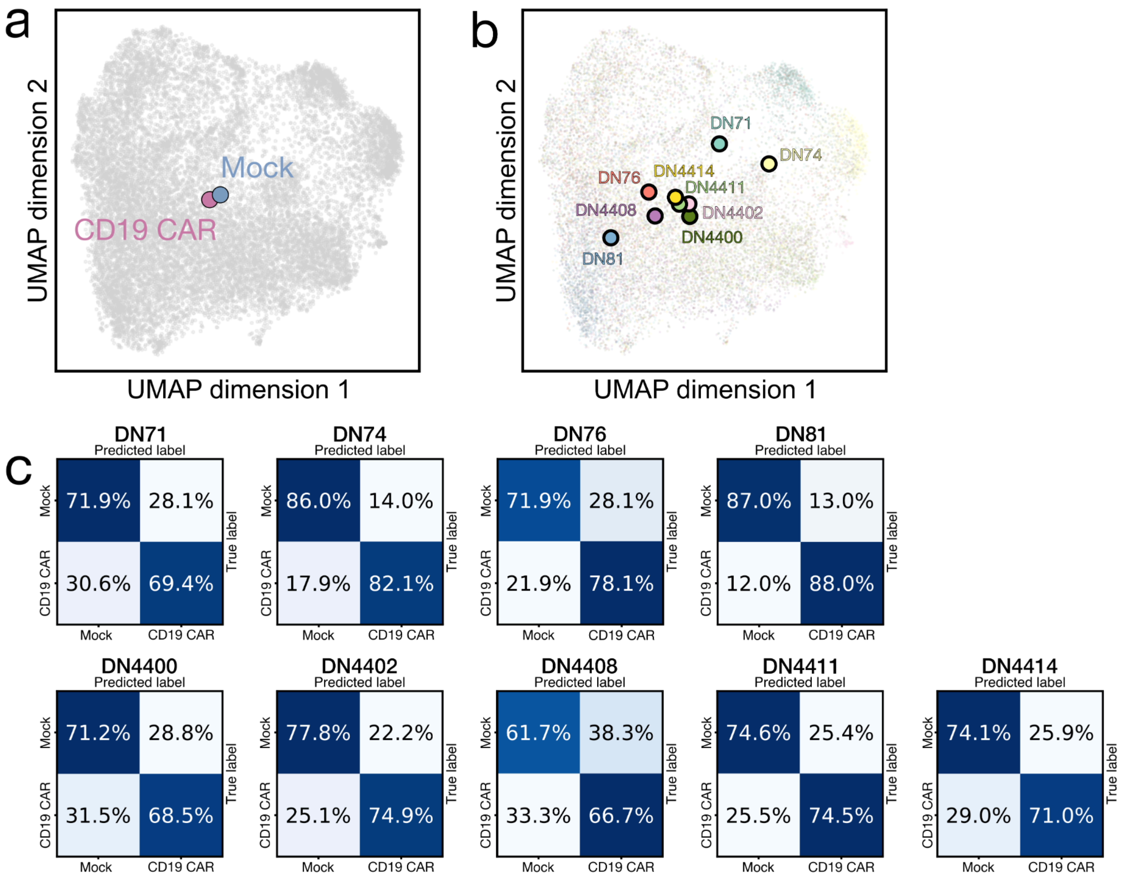


**Supplementary Fig. 6:** Pre-sorted CD19-CAR spectral analysis across donors

**(a-b)** Two-dimensional Uniform Manifold Approximation and Projections (UMAPs) of CD19-CAR and Mock Raman spectra from nine donors, generated after Principal Component Analysis (PCA) reduction and balanced by class and donor. **(a)** Class centroids and **(b)** donor centroids are overlaid, illustrating that inter-donor variability exceeds between-class separations. This strong donor prior motivated a per-donor model training for our dataset.[14] **(c)** Normalized confusion matrices for each donor showing classification accuracies for CD19-CAR and Mock spectra using a LightGBM classifier with 10-fold stratified cross-validation.

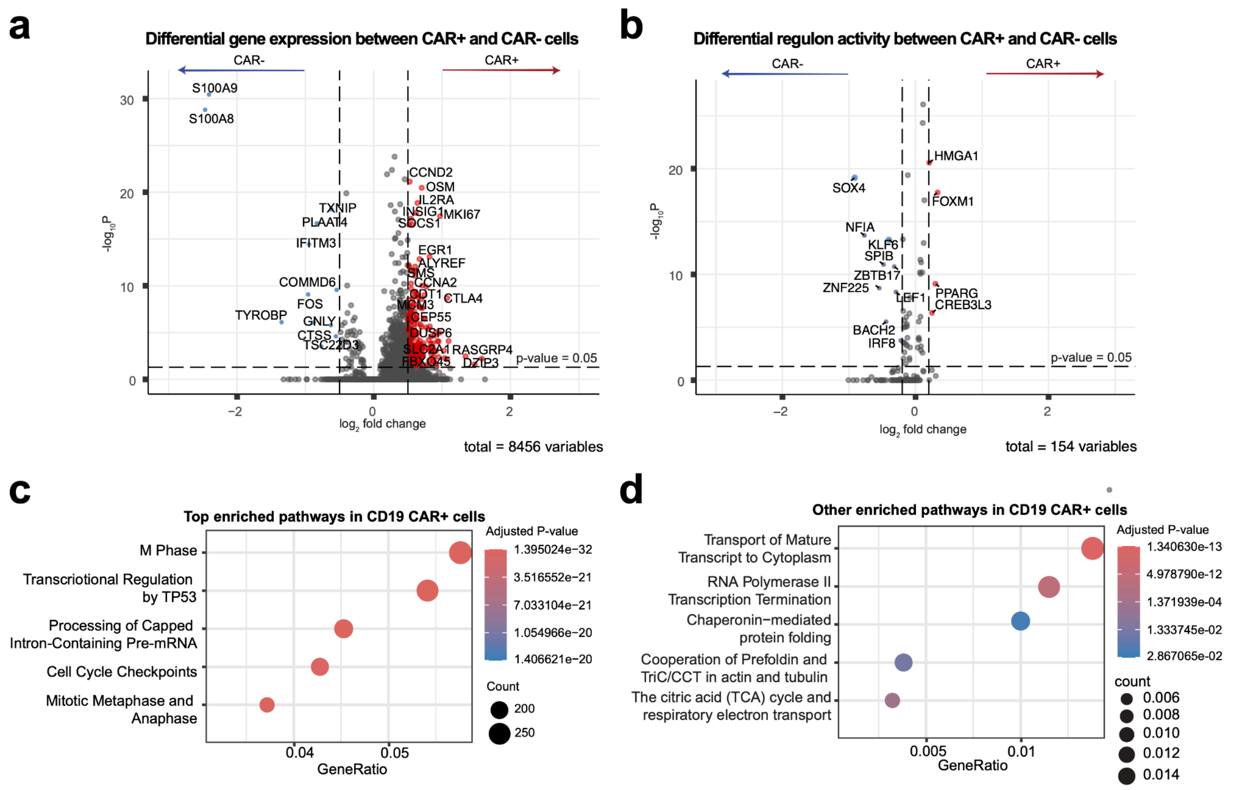


**Supplementary Fig. 7:** Transcriptomic signatures of CAR+ versus CAR- cells in CD19 CAR T infusion products

**(a)** Differential gene expression (DEG) volcano plot comparing CAR+ and CAR- infusion products. DEGs were selected based on log2 fold change |log2FC| > 0.3 and p-value < 0.05. **(b)** Differential regulon activity volcano plot comparing CAR+ and CAR- infusion products. **(c)** Dot plot showing top enriched pathways in CAR+ cells. **(d)** Additional enriched pathways in CAR+ groups, related to protein folding and aromatic expression.

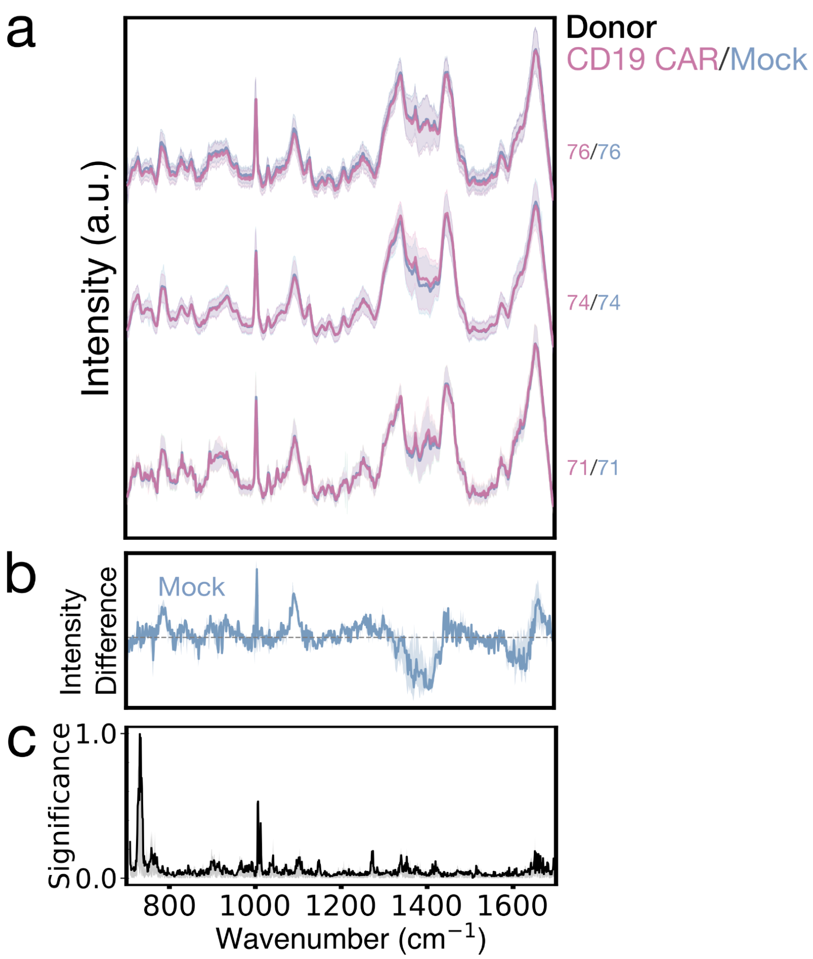


**Supplementary Fig. 8:** Non-SERS CD19 CAR vs Mock Raman data

**(a)** Waterfall plot of mean normalized spectra of live single CD19-CAR (pink) and Mock (blue) T cells with ±1 SD (shaded), separated by donor (identifier labeled on the right). **(b)** Median donor-normalized spectral difference of Mock cells relative to CAR with 25-75% quantile range. The grey dashed line marks zero difference. **(c)** Mean feature importance plot depicting the contribution of each spectral dimension (wavenumber) to classification using a trained LightGBM model.

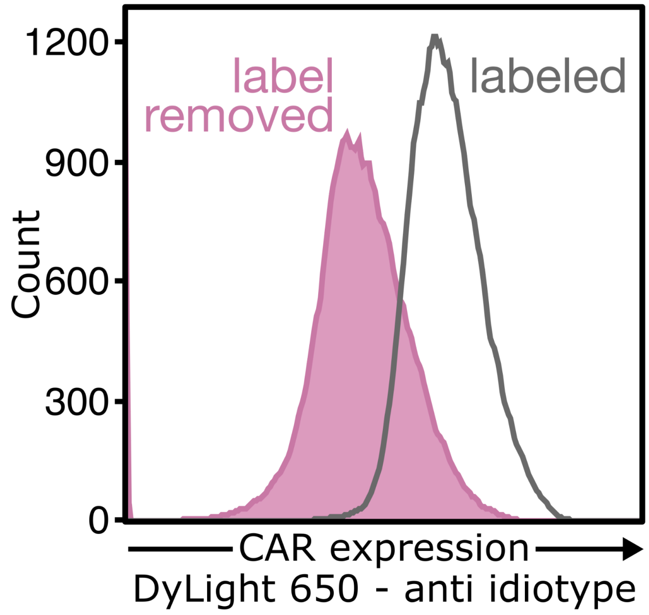


**Supplementary Fig. 9:** Loss of anti-idiotype label after post-sort incubation

Flow cytometry of purified CD19-CAR T cells following overnight incubation and washing. Cells analyzed without re-staining ("label removed", pink) show a substantial reduction in fluorescent signal compared to re-labeled cells with the same antibody ("labeled", grey). The decline in fluorescence post-incubation is consistent with loss of the initially bound antibody, potentially due to dissociation or internalization and degradation of the receptor-ligand complex. Re-labeling serves as a positive control confirming that CAR expression was maintained and the loss of signal in the "label removed" condition was due to anti-idiotype removal.

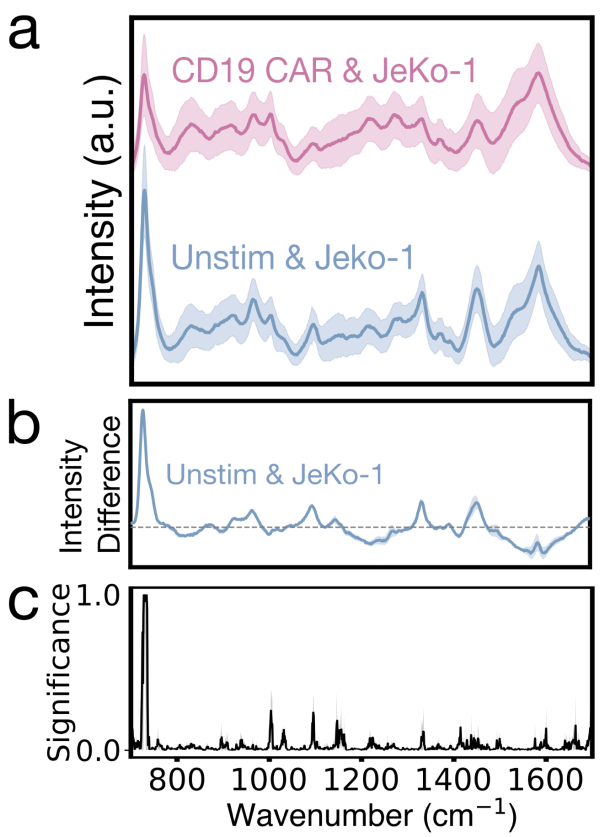


**Supplementary Fig. 10:** Population-level co-culture Raman signatures

**(a)** Mean normalized SERS spectra of live CD19-CAR (pink) and unstimulated (Unstim; blue) T cells co-cultured with JeKo-1 B cells, aggregated over the full 1.5-hour recording period (two donors), with ±1 SD (shaded). **(b)** Median donor-normalized spectral difference of Unstim cells relative to CAR T cells, with 25-75% quantile range. The grey dashed line marks zero difference. **(c)** Mean feature importance plot depicting the contribution of each spectral dimension (wavenumber) to classification using a trained LightGBM model.

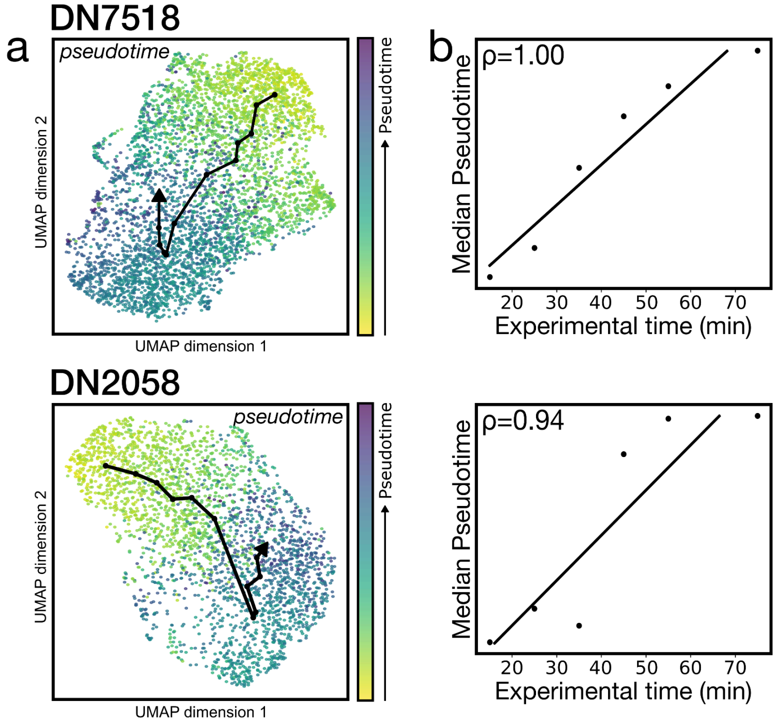


**Supplementary Fig. 11:** Pseudotime trajectory analysis and correlation with real time

**(a)** Pseudotime trajectories overlaid on UMAPs for two donors by pseudotime color scale (yellow to purple). The arrow indicates the trajectory backbone, connecting the centroids of pseudotime bins. **(b)** Comparison of experimental time and pseudotime across binned data revealed a strong correlation (Spearman's rank correlation: $\rho = 1.00$ for DN7518, $p < 0.001$, and $\rho = 0.94$ for DN2058, $p = 0.0048$). The linear regression trendline is shown for visualization; correlation was assessed using Spearman's rank correlation. Deviations from perfect monotonicity may be due to the activation heterogeneity across cells.

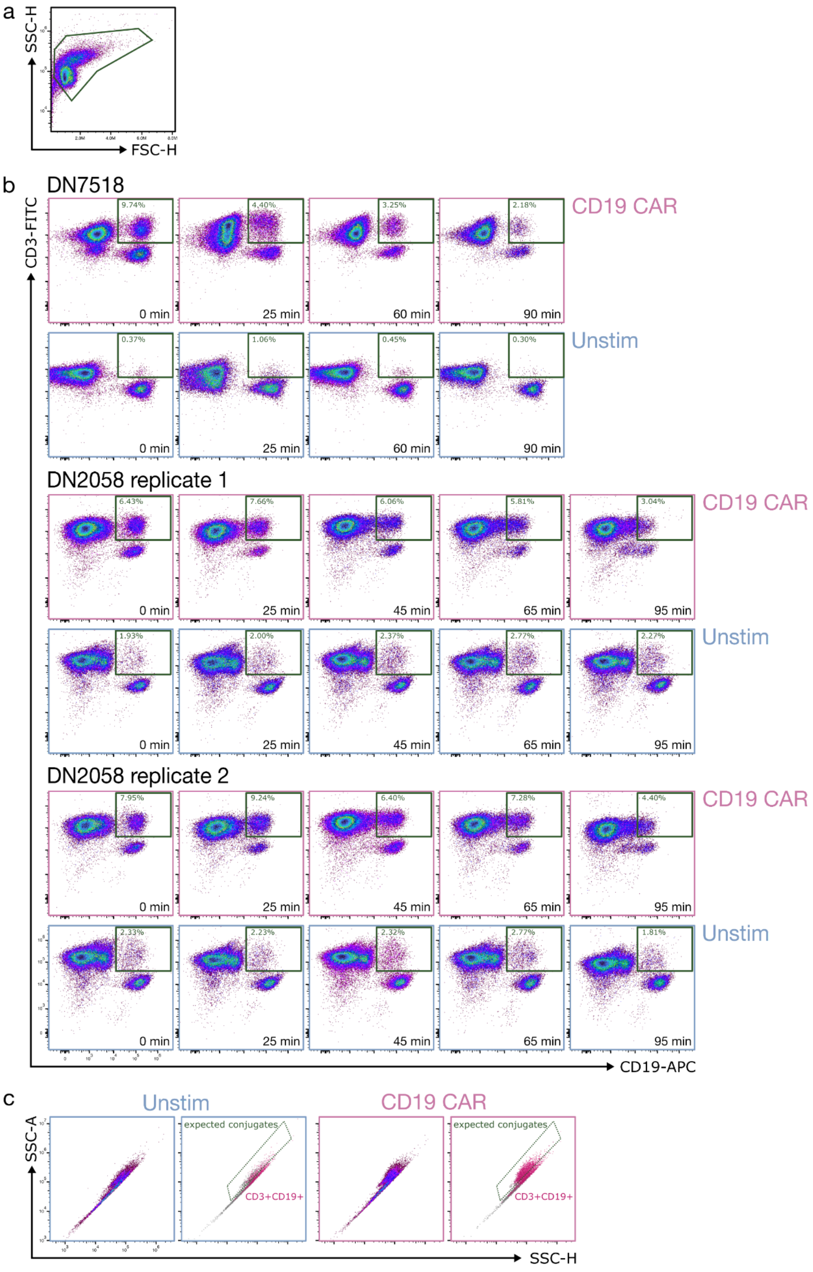


**Supplementary Fig. 12:** Flow cytometry characterization of antigen-specific activation

Flow cytometry density plots for CD19-CAR (pink) and unstimulated (Unstim; blue) T cells co-cultured with JeKo-1 B cells for approximately 90 minutes across two donors. **(a)** Density flow plot showing the debris-removal gate (FSC-H vs SSC-H). No singlet/doublet gate was applied to preserve T-B cell conjugates in the CD3+CD19+ events. **(b)** CD3 vs CD19 density plots for all donors, replicates, and conditions across all measured time points, showing the strong presence of the CD3+CD19+ population (green gate) in the CD19-CAR condition, which is absent in the Unstim controls. **(c)** SSC-H vs SSC-A distributions for all events after debris removal in Unstim and CD19-CAR co-cultures at a representative timepoint (25 min) (left). CD3+CD19+ events overlaid (pink) on the full event distribution (grey). In the CD19-CAR condition, these events localize to a high-side-scatter region ("expected conjugates"), consistent with interacting cells, whereas Unstim samples show few such events.

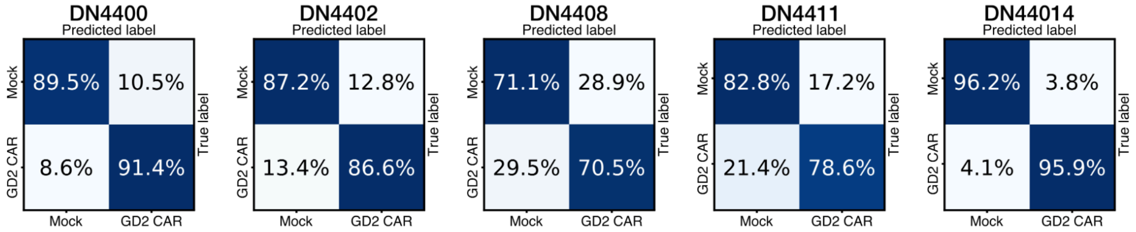


**Supplementary Fig. 13:** GD2-CAR and Mock SERS spectral classification across donors

Normalized confusion matrices for five donors showing classification accuracies for GD2-CAR and Mock spectra using a LightGBM classifier with 10-fold stratified cross-validation.

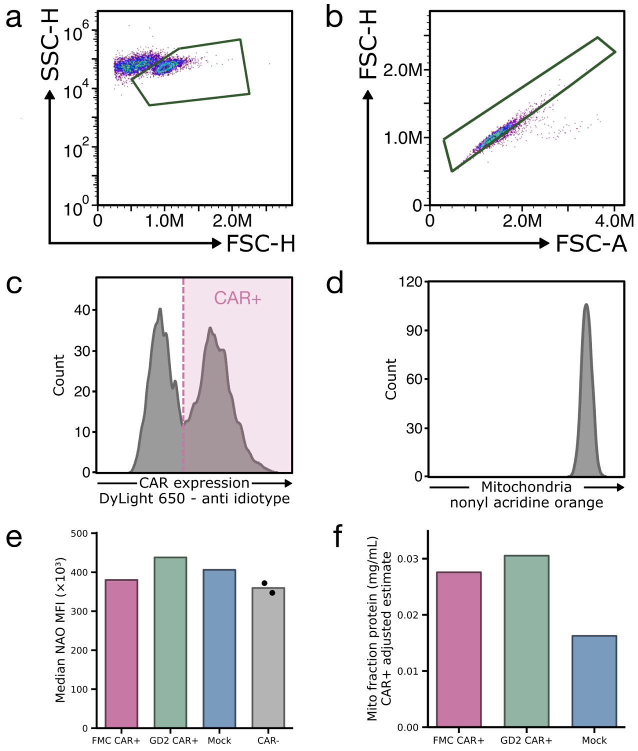


**Supplementary Fig. 14:** Representative mitochondrial measurements in FMC-CAR, GD2-CAR, and Mock T cells

Flow cytometry gating strategy for mitochondrial mass quantification. Flow cytometry plots show gating for **(a)** lymphocytes, **(b)** singlets, and **(c)** CAR-expressing cells with CAR+ gate indicated (anti-CAR staining; DyLight 650). **(d)** Mitochondrial mass was assessed using nonyl acridine orange (NAO). **(e)** Median NAO fluorescence intensity (MFI) is plotted for CAR-positive FMC-CAR and GD2-CAR T cell populations, the full Mock T cell population, and CAR-negative cells (negative gate) from the CAR-transduced samples (representative donor). **(f)** Representative mitochondrial extraction followed by Bradford assay quantification of mitochondrial protein concentration. For CAR-transduced samples, concentrations were adjusted by the CAR+ transduction efficiency measured using flow cytometry to estimate the protein concentration per CAR+ cell. Mock was unadjusted.

**Supplementary Table 5:** Expanded flow cytometry and CyTOF validation values for longitudinal patient samples

| CAR T cell frequency: % CAR+ of CD3+ parent population | | | | |
|---|---|---|---|---|
| **Patient ID** | **Timepoint** | **Fresh flow (%)** | **Thawed CyTOF (%)** | **Thawed flow (%)** |
| 016 | Peak | 11.4 | 18.0 | 22.8 |
| | Late | 2.9 | 0.08 | 0.2 |
| 019 | Peak | 21.1 | 6.15 | 12.3 |
| | Late | 22.9 | 25.7 | 25.2 |
| 022 | Peak | 34.6 | 22.9 | 47.5 |
| | Late | 20.0 | 17.1 | 24.3 |
| 024 | Peak | 19.5 | 13.9 | 19.5 |
| | Late | 2.9 | 3.9 | 5.2 |

| CD4/CD8 composition and proliferation: % of CD3+ parent population | | | | |
|---|---|---|---|---|
| **Patient ID** | **Timepoint** | **CD4+ (%)** | **CD8+ (%)** | **Ki-67+ (%)** |
| 016 | Pre | 18.3 | 66.3 | 6.5 |
| | Peak | 8.4 | 76.6 | 70.1 |
| | Late | 7.2 | 89.8 | 26.1 |
| 019 | Pre | 6.4 | 58.8 | 4.4 |
| | Peak | 3.8 | 31.0 | 26.3 |
| | Late | 7.1 | 65.3 | 19.0 |
| 022 | Pre | 23.1 | 0.3 | 5.9 |
| | Peak | 8.3 | 0.3 | 13.3 |

| | | | | |
|---|---|---|---|---|
| | Late | 25.5 | 0.4 | 39.1 |
| 024 | Pre | 11.2 | 0.9 | 2.1 |
| | Peak | 4.9 | 0.6 | 33.1 |
| | Late | 6.6 | 1.1 | 27.7 |

**CD57/T-bet differentiation-associated phenotypes: % of CD3+ parent population**

| Patient ID | Timepoint | CD57+ T-bet+ (%) | CD57- T-bet+ (%) | CD57- T-bet- (%) |
|---|---|---|---|---|
| 016 | Pre | 39.4 | 14.9 | 37.3 |
| | Peak | 37.4 | 38.0 | 17.9 |
| | Late | 48.4 | 39.4 | 7.7 |
| 019 | Pre | 42.4 | 7.7 | 40.3 |
| | Peak | 24.1 | 4.4 | 65.7 |
| | Late | 53.0 | 16.7 | 24.7 |
| 022 | Pre | 23.2 | 36.2 | 36.7 |
| | Peak | 10.4 | 23.1 | 62.6 |
| | Late | 22.2 | 53.0 | 22.5 |
| 024 | Pre | 63.1 | 9.9 | 13.3 |
| | Peak | 40.5 | 12.8 | 39.3 |
| | Late | 58.8 | 24.1 | 13.1 |

**CD39/PD-1 phenotypes: % of CD3+ parent population**

| Patient ID | Timepoint | CD39+ PD-1+ (%) | CD39- PD-1+ (%) | CD39- PD-1- (%) |
|---|---|---|---|---|
| | Pre | 5.3 | 41.2 | 45.0 |

| | | | | |
|---|---|---|---|---|
| 016 | Peak | 10.4 | 39.7 | 35.1 |
| | Late | 0.9 | 31.1 | 66.2 |
| 019 | Pre | 0.7 | 12.9 | 56.3 |
| | Peak | 1.8 | 12.2 | 23.6 |
| | Late | 1.1 | 35.7 | 42.0 |
| 022 | Pre | 2.8 | 8.6 | 2.8 |
| | Peak | 19.4 | 7.7 | 19.4 |
| | Late | 8.0 | 22.2 | 8.0 |
| 024 | Pre | 1.2 | 2.2 | 89.6 |
| | Peak | 11.3 | 3.9 | 37.4 |
| | Late | 6.9 | 8.9 | 61.2 |

**Note**: “Fresh flow” refers to flow cytometry performed on fresh clinical samples shortly after patient blood collection. “Thawed CyTOF” refers to matched CyTOF data generated from thawed aliquots in a prior study.[15] “Thawed flow” refers to matched flow cytometry data acquired immediately before Raman spectral acquisition on the same samples used for SERS analysis. CAR T cell frequency values are reported from all three sources. Because thawed flow was acquired from the same samples immediately before Raman analysis, it was used as the most sample-matched reference for CAR+ T cell frequency in Fig. 6f. All other immune state measurements in this table and in Fig. 6b were acquired by CyTOF on matched thawed samples. Longitudinal trends were generally consistent across assays. Differences in CAR+ T cell frequencies across fresh flow, thawed CyTOF, and thawed flow likely reflect differences in assay platform and sample preparation, including cryopreservation and thawing. Because the Raman analyses were not intended to provide calibrated CAR+ T cell quantification, these measurements were used as complementary orthogonal validation.

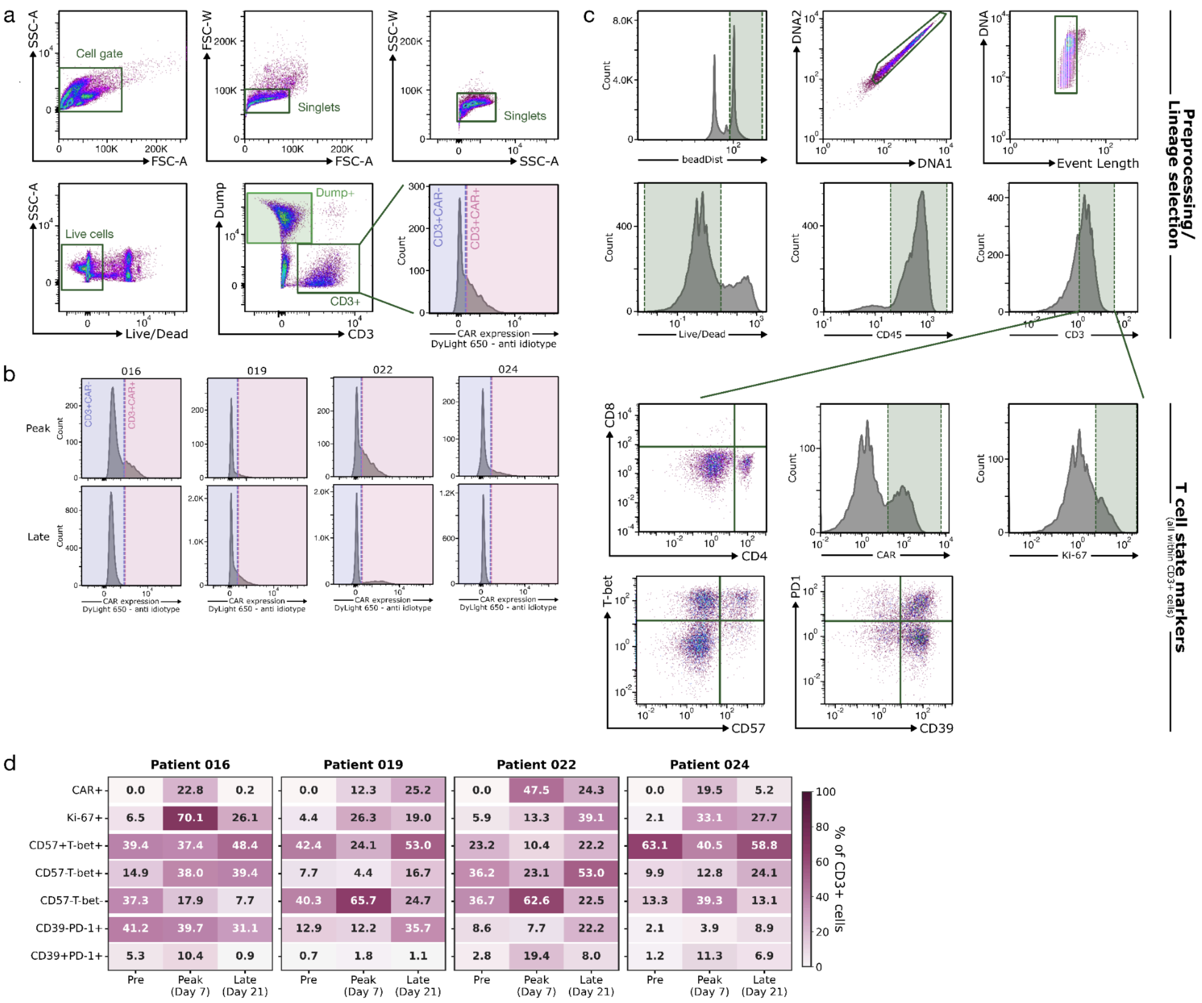


**Supplementary Fig. 15:** Representative gating for fluorescence-activated cell sorting (FACS) and mass cytometry (CyTOF) analysis of longitudinal patient samples

**(a)** Representative FACS gating strategy used to isolate PBMC populations for downstream Raman analysis. Flow cytometry plots show gates used to select intact cells (upper left), singlets (upper middle and right), and live cells (lower left). Dump+ cells (light green) were identified using a combined lineage Dump channel containing CD19+, CD14+, and CD235a+ events (lower middle). From the CD3+ parent population, CD3+ CAR+ (pink) and CD3+ CAR- (blue) populations were isolated (lower right). **(b)** Final CAR expression gates for Peak and Late timepoints across all patients. **(c)** Representative CyTOF gating strategy used to analyze matched patient and timepoint data from a prior study.[15] Full cohort data are provided in Supplementary Table 5. Preprocessing and lineage selection plots show gates used to select cells (upper left), singlets (upper middle and right), live cells (lower left), leukocytes (CD45+; lower middle), and T cells (CD3+; lower right). From the CD3+ parent population, cells were analyzed for various T cell state markers, including T cell lineage composition (CD4 vs CD8; upper left), CAR T cell frequency (CAR+; upper middle), proliferation (Ki-67+; upper right), differentiation-associated phenotypes (CD57 vs T-bet; lower left), and inhibitory or exhaustion-associated phenotypes (CD39 vs PD1; lower middle). **(d)** Heatmap summarizing patient- and timepoint-specific immune cell states, including CAR T cell frequency measured by flow cytometry and Ki-67, CD57/T-bet, and CD39/PD-1 phenotypes measured by CyTOF. All populations are reported as a percentage of CD3+ cells.

**Supplementary Table 6:** Spectral count for longitudinal patient samples (Figure 6)

| Timepoint | Sample Type | Spectra count |
|---|---|---|
| **Patient ID 016** | | |
| Pre | Mixed PBMCs | 923 |
| | Sorted CD3+CAR- | 649 |
| | Sorted Dump+ | 245 |
| Peak | Mixed PBMCs | 382* |
| | Sorted CD3+CAR- | 280 |
| | Sorted CD3+CAR+ | 37 |
| | Sorted Dump+ | 263 |
| Late | Mixed PBMCs | 1069 |
| **Patient ID 019** | | |
| Pre | Mixed PBMCs | 596 |
| | Sorted CD3+CAR- | 695 |
| | Sorted Dump+ | 594 |
| Peak | Mixed PBMCs | 596 |
| | Sorted CD3+CAR- | 74 |
| | Sorted CD3+CAR+ | 39 |
| | Sorted Dump+ | 600 |
| Late | Mixed PBMCs | 703 |
| | Sorted CD3+CAR- | 704 |
| | Sorted CD3+CAR+ | 711 |
| | Sorted Dump+ | 45 |
| **Patient ID 022** | | |
| | Mixed PBMCs | 644 |

| | | |
|---|---|---|
| Pre | Sorted CD3+CAR- | 680* |
| | Sorted Dump+ | 535 |
| Peak | Mixed PBMCs | 531 |
| | Sorted CD3+CAR- | 387 |
| | Sorted CD3+CAR+ | 561 |
| | Sorted Dump+ | 411 |
| Late | Mixed PBMCs | 631 |
| | Sorted CD3+CAR- | 626* |
| | Sorted CD3+CAR+ | 565 |
| **Patient ID 024** | | |
| Pre | Mixed PBMCs | 847 |
| | Sorted CD3+CAR- | 755 |
| | Sorted Dump+ | 640* |
| Peak | Mixed PBMCs | 495 |
| | Sorted CD3+CAR- | 530 |
| | Sorted CD3+CAR+ | 75 |
| | Sorted Dump+ | 464 |
| Late | Mixed PBMCs | 750 |
| | Sorted CD3+CAR- | 751 |
| | Sorted CD3+CAR+ | 313 |
| | Sorted Dump+ | 623 |

**Note**: Spectral counts are of analyzed spectra post-filtering for laser-induced damage or detector saturation (Supplementary Fig. 4). Dump+ sorted population consisted of CD19+, CD14+, and CD235a+ cells.

* Spectra indicated with an asterisk were noted during acquisition to partially overlap the edge of the liquid well. Off-well spectra were excluded based on the procedure noted in Methods.

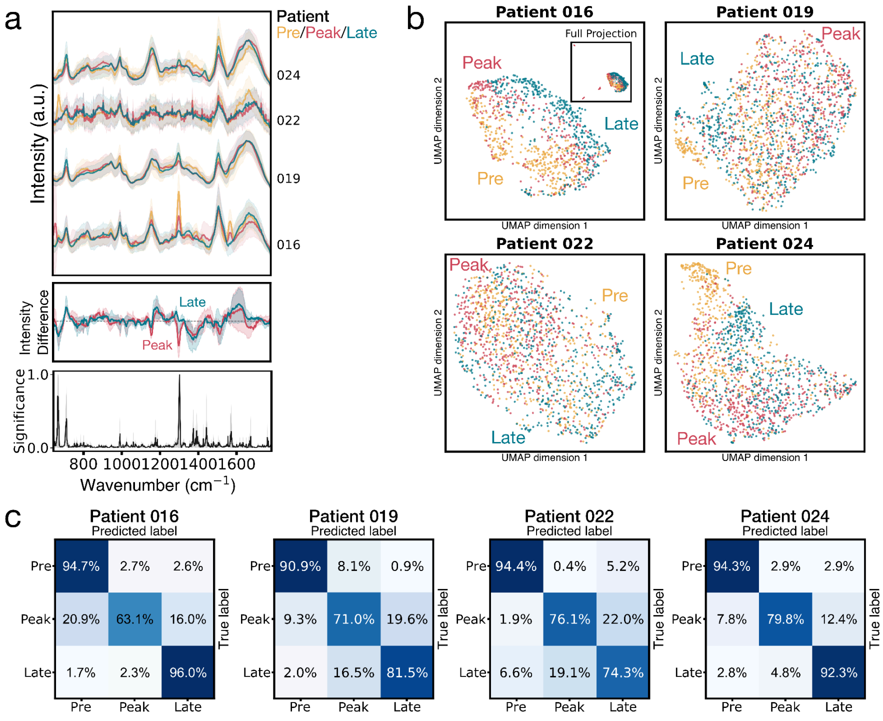


**Supplementary Fig. 16:** Longitudinal mixed PBMC clinical sample analysis across patients

**(a)** Waterfall plot of mean normalized SERS spectra of live patient PBMCs at Pre (yellow), Peak (red), and Late (blue-green) timepoints with ±1 SD (shaded), separated by patient (identifier labeled on the right) (top). Median patient-normalized spectral differences for Peak and Late PBMC spectra relative to the matched Pre timepoint are shown with 25-75% quantile ranges (middle). The grey dashed line marks zero difference. Mean feature importance plot showing the contribution of each spectral dimension (wavenumber) to multiclass timepoint classification across trained patient-specific LightGBM models (bottom). **(b)** Two-dimensional UMAP embeddings of mixed PBMC Raman spectra for each patient, generated after PCA reduction and colored by timepoint. **(c)** Normalized confusion matrices for each patient showing multiclass timepoint classification accuracies for Pre, Peak, and Late mixed PBMC spectra using a LightGBM classifier with 10-fold stratified cross-validation. These analyses support the data displayed in Fig. 6b-c.

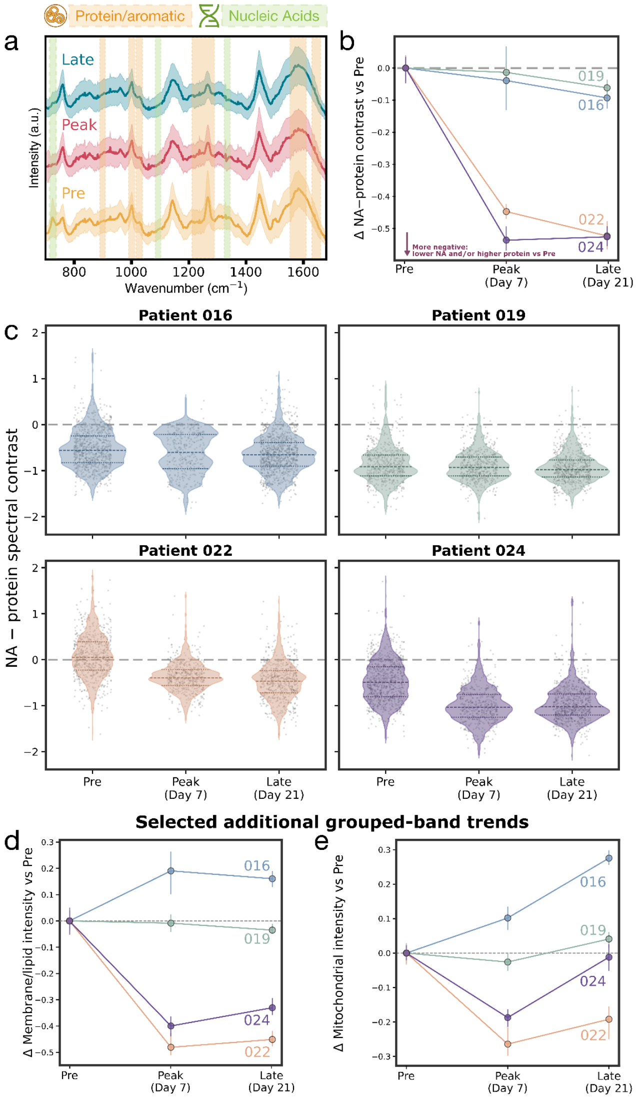


**Supplementary Fig. 17:** Longitudinal PBMC biochemical band intensity trends

**(a)** Waterfall plot of mean normalized SERS spectra of live patient PBMCs across four patients at Pre (yellow), Peak (red), and Late (blue-green) timepoints with ±1 SD (shaded). Colored spectral bands highlight protein/aromatic- (gold) and nucleic acid- (green) associated regions for grouped band analysis. **(b)** Longitudinal PBMC Raman intensity trends showing the difference between averaged nucleic acid (NA)-associated and protein/aromatic-associated bands relative to each patient's matched Pre sample. Points show medians; error bars indicate 95% bootstrap confidence intervals. Negative values indicate lower NA-associated contribution and/or higher protein/aromatic-associated signal relative to matched Pre spectra. **(c)** Single-spectrum distributions of the difference between combined nucleic acid-associated and protein/aromatic-associated band intensities for each patient and timepoint. Individual points represent single spectra. Dashed internal lines indicate the median and interquartile range. **(d)** Selected additional grouped-band Raman trends calculated from membrane/lipid-associated regions at 717–721, 870–880, and 1455–1465 $cm^{-1}$ (left)[4,6] and mitochondrial-associated regions at 749–755, 1130–1131, 1310–1311, and 1584–1590 $cm^{-1}$ (right),[9] shown relative to each patient's matched Pre sample (patients differentiated by color). Band assignments are approximate and may include overlapping molecular contributions.

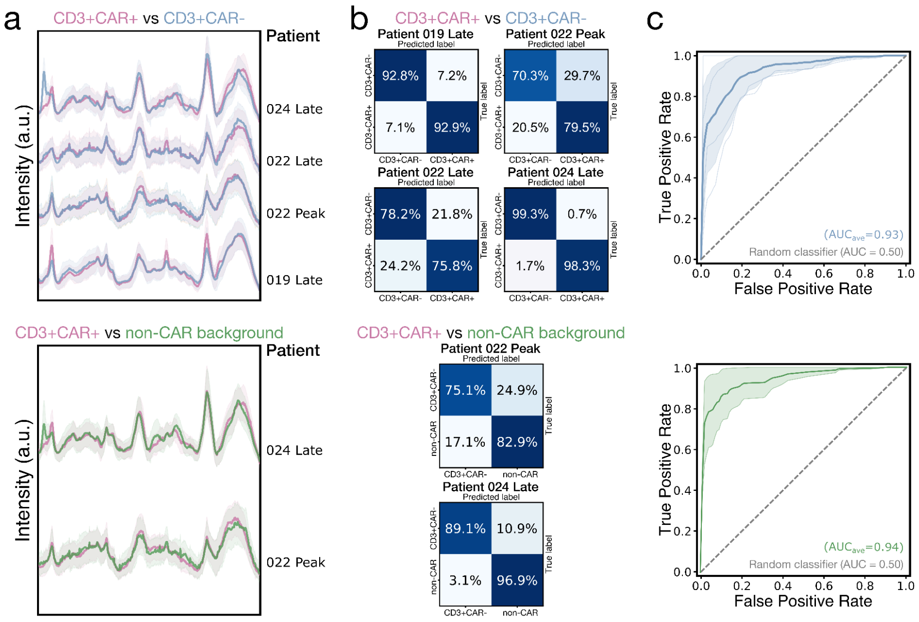


**Supplementary Fig. 18:** Longitudinal flow-isolated clinical sample analysis across patients included in Fig. 6e

**(a)** Waterfall plot of mean normalized SERS spectra from flow-isolated CD3+CAR+ (pink) and CD3+CAR- (blue) T cells (top) and CD3+CAR+ versus non-CAR background cells (green) (bottom), with ±1 SD (shaded). Spectra are separated by patient/timepoint grouping with sufficient spectral counts (identifier labeled on the right). **(b)** Normalized confusion matrices for each patient/timepoint grouping showing binary classification performance for CD3+CAR+ versus CD3+CAR- T cell spectra (top) and CD3+CAR+ versus non-CAR background cell spectra (bottom) using a LightGBM classifier with 10-fold stratified cross-validation. **(c)** ROC curves displaying classification performance of CD3+CAR+ versus CD3+CAR- T cell spectra (top) and CD3+CAR+ versus non-CAR background spectra (bottom). Thick lines show the mean across all patient/timepoint comparisons, shaded regions indicate ±1 SD, and thin lines show individual patient/timepoint curves. The grey dashed line represents a random classifier (AUC = 0.5). Mean AUCs are given in each plot. Non-CAR background consisted of both Dump+ and CD3+CAR- populations, as described in Methods. These analyses support the data displayed in Fig. 6e.

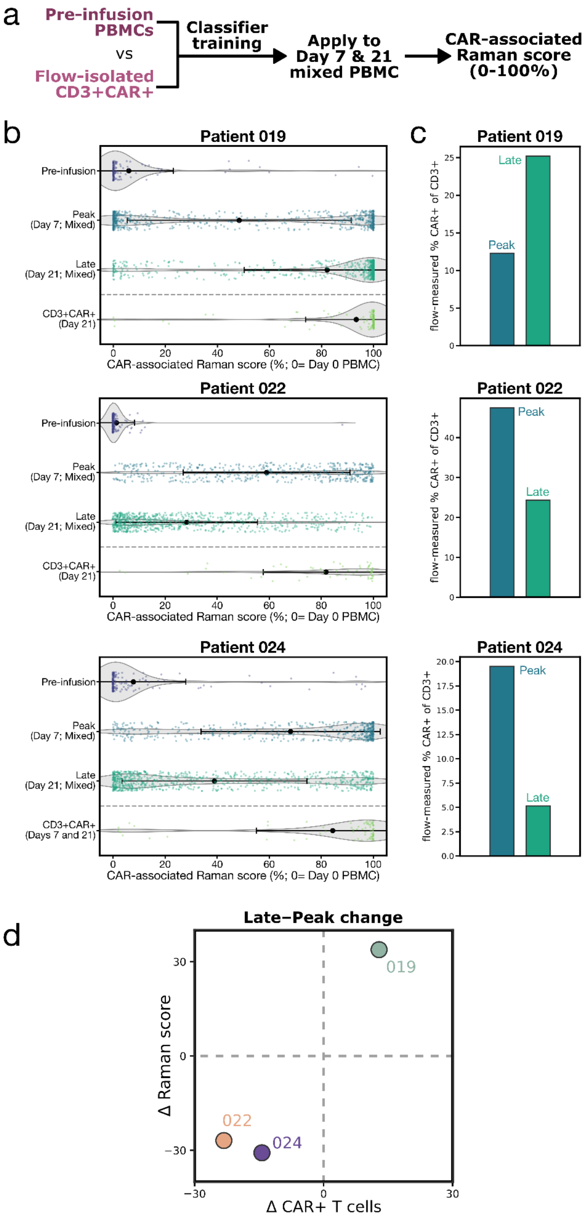


**Supplementary Fig. 19:** Clinical sample CAR-associated Raman score for patients included in Fig. 6f

**(a)** Diagram defining the CAR-associated Raman score. A classifier was trained to distinguish pre-infusion mixed PBMC spectra from flow-isolated CD3+CAR+ spectra from post-infusion timepoints with sufficient spectral counts (Methods). For Patient 022, Peak and Late

CD3+CAR+ spectra were pooled. For Patients 019 and 024, only Late CD3+CAR+ spectra were used because of low Peak spectral counts. The trained classifier was then applied separately to Peak and Late mixed PBMC spectra. Patient 016 was excluded from this analysis because no post-infusion CD3+CAR+ sample met the inclusion threshold. The classifier output probability was defined as the "CAR-associated Raman score". **(b)** Single-spectrum distributions of CAR-associated Raman scores for held-out pre-infusion PBMC and flow-isolated CD3+CAR+ test spectra, as well as Peak and Late mixed PBMC spectra. Points represent individual spectra; black markers and horizontal bars indicate the mean ±SD. **(c)** Matched flow-measured CAR+ T cell frequencies within the CD3+ parent population for Peak and Late timepoints. These measurements illustrate the temporal direction of CAR+ T cell frequency for comparison with the Raman score distributions in (b). **(d)** Late-Peak change plot comparing changes in CAR-associated Raman score and flow-measured CAR+ T cell frequency from Peak to Late. Points in the first and third quadrants indicate concordant directional changes. These analyses support the data displayed in Fig. 6f.
